# Cell-specialized chloroplast signaling orchestrates photosynthetic and extracellular reactive oxygen species for stress responses

**DOI:** 10.1101/2023.08.02.551742

**Authors:** Estee E. Tee, Stephen J. Fairweather, Hanh M. Vo, Chenchen Zhao, Andrew Breakspear, Sachie Kimura, Melanie Carmody, Michael Wrzaczek, Stefan Bröer, Christine Faulkner, Jaakko Kangasjärvi, Zhong-Hua Chen, Barry J. Pogson, Kai Xun Chan

## Abstract

Cellular responses to abiotic stress involve multiple signals including secondary messengers such as reactive oxygen species (ROS) and Ca^2+^, phytohormones such as abscisic acid (ABA) and chloroplast-to-nucleus retrograde signals such as 3’-phosphoadenosine 5’-phosphate (PAP). Mechanism(s) by which these messengers, produced in different subcellular compartments, intersect for cell regulation remain enigmatic. Previously we showed that the chloroplast retrograde signal PAP, similar to ABA, induces an increase in ROS levels in guard cells (Pornsiriwong *et al*, 2017). Here we demonstrate a mechanistic link enabling ABA and PAP to coordinate both chloroplast and plasma membrane ROS production. In whole leaves, PAP alters various ROS-related processes including plasmodesmal permeability as well as responses to ozone and the bacterial elicitor flg22, but mainly initiates processes that quench ROS during oxidative stress. Conversely, we show in guard cells, both PAP and ABA induce an increase in ROS levels in both chloroplasts via photosynthetic electron transport, and the apoplast via the RESPIRATORY BURST OXIDASE HOMOLOG (RBOH). Both subcellular ROS sources were necessary for ABA- and PAP-mediated stomatal closure. However, PAP signaling diverges from ABA by activating RBOHD, instead of RBOHF, for apoplastic ROS mediated stomatal closure. We identified three calcium-dependent protein kinases (CPKs) as the post-translational activators of RBOHD-mediated ROS production. CPK13, CPK32, and CPK34 were transcriptionally induced by PAP and concurrently activate RBOHD and the slow anion channel SLAC1 by phosphorylating two Serine (S) residues, including S120 which is also targeted by the core ABA signaling kinase OPEN STOMATA 1 (OST1). Consequently, overexpression of the PAP-induced CPKs rescues stomatal closure in *ost1.* Our data identify stomatal chloroplasts, to be nodes in the multifaceted cellular stress response networks as they are both sources and mediators of ROS and retrograde signals such PAP. Thus, chloroplasts are not just mediators of photosynthesis in response to, for example, excess light, but can serve as critical nodes in the multifaceted cellular stress response networks in specialized cells via retrograde signals, providing support to the concept of sensory plastids.

**Significance Statement:** The chloroplast is an environmental sensor for stresses such as excess light and drought via the activation of photosynthetic-mediated retrograde signals. However, how does it function in specialized cells for which carbon fixation is secondary? Here we show the chloroplast is an important node to coordinate multiple plant signaling pathways in response to stresses such as drought. The chloroplast retrograde signal 3’-phosphoadenosine 5’-phosphate (PAP) plays multiple roles in reactive oxygen species (ROS) signaling and homeostasis. While PAP suppresses ROS in photosynthetic tissue, PAP instead induces guard cell ROS in chloroplasts and extracellular space to induce stomatal closure. We decipher how PAP-induced proteins activate both extracellular ROS production and anion channels for stomatal closure, thus providing a mechanism by which chloroplasts provide a strategic complement to canonical hormonal pathways in regulating plant physiological responses in specialized cells.

## Introduction

Plant responses to abiotic and biotic stresses involve a suite of coordinated cellular signaling cascades. These involve multiple messenger molecules including reactive oxygen species (ROS), calcium ions (Ca^2+^), phytohormones such as abscisic acid (ABA), and retrograde signals, defined as signals from an organelle (i.e., the chloroplast or mitochondria) that regulate nuclear gene expression (Chan et al., 2016b; Cutler et al., 2010; Waszczak et al., 2018). While each of the aforementioned messengers have been extensively studied, their spatio-temporal intersections remain enigmatic. For example, both apoplastic ROS and organelle-to-nucleus retrograde signals occur concurrently, yet it is unclear how these two classes of signals interact and are coordinated in specialized cells. This is relevant to the proposition that plants have specialized plastids for sensing and signaling (Mackenzie and Mullineaux, 2022).

ROS are crucial signaling molecules produced in diverse subcellular compartments including chloroplasts and the apoplast. Several types of ROS exist in plant cells and distinct ROS types play different signaling roles (Chan et al., 2016b; Cutler et al., 2010; Waszczak et al., 2018), but herein we refer to chloroplast and apoplast “ROS” as an encompassing term for three ROS types formed via similar mechanisms at these subcellular locations: superoxide (O_2_^-^), its dismutation product hydrogen peroxide (H_2_O_2_), and hydroxyl radical (OHꞏ); with H_2_O_2_ being the most stable (Waszczak et al., 2018). ROS generated from photosystem I (PSI) in chloroplasts participates in chloroplast-to-nucleus retrograde signaling, both by triggering production of retrograde signals via H_2_O_2_-responsive redox regulation (Chan et al., 2016a) and by itself acting as a retrograde signal (Exposito-Rodriguez et al., 2017). Silencing the chloroplastic H_2_O_2_ scavenging enzyme, thylakoid membrane-bound ascorbate peroxidase (tAPX), increased chloroplastic H_2_O_2_ leading to the up-regulation of multiple stress-associated genes, including pathogen-responsive genes in photosynthetic tissues (Maruta et al., 2012). High light stress increases H_2_O_2_ production in leaf chloroplasts; the chloroplasts in close physical association with the nucleus can transfer their H_2_O_2_ into the nucleus, potentially representing a direct means for H_2_O_2_ to act as a retrograde signal to induce nuclear gene expression (Exposito-Rodriguez et al., 2017). However, the nuclear targets of chloroplastic-sourced H_2_O_2_ are still unknown.

In contrast to chloroplast PSI ROS, apoplastic superoxide and H_2_O_2_ are well known to act as secondary messengers in dynamic signaling cascades. One such well characterized pathway that utilizes apoplastic and cellular ROS as a secondary messenger is abscisic acid (ABA)-mediated stomatal closure. In response to an accumulation of ABA under stress conditions such as drought, a simplified guard cell process occurs as follows: ABA forms a complex with PYRABACTIN RESISTANCE 1/PYR1-LIKE/REGULATORY COMPONENTS OF ABA RECEPTORS, with ABSCISIC ACID-INSENSITIVE 1 (ABI1) PP2C phosphatase then binding to the complex (Ma et al., 2009; Park et al., 2009). This removes the inhibition of SUCROSE NON-FERMENTING-1 RELATED PROTEIN KINASE (SnRK) 2.6 (SnRK 2.6)/SRK2E/OPEN STOMATA 1 (OST1), which can auto-phosphorylate (Belin et al., 2006; Yoshida et al., 2006). OST1 interacts with RESPIRATORY BURST OXIDASE HOMOLOG F (RBOHF) and RBOHD, leading to RBOH-mediated ROS (superoxide and H_2_O_2_) production (Acharya et al., 2013; Sirichandra et al., 2009). OST1 also directly regulates the activity of multiple ion channels by phosphorylation, including activation of SLOW ANION CHANNEL 1 (SLAC1) and rapid type anion channel ALMT12 (Acharya et al., 2013; Geiger et al., 2009; Imes et al., 2013), and inhibiting inward rectifying POTASSIUM CHANNEL (KAT1) (Sato et al., 2009). While OST1 has the capacity to directly act on the channels, both ROS and cytosolic Ca^2+^ are required for stomatal closure with H_2_O_2_ activating Ca^2+^ channels to induce the influx of Ca^2+^ and release of Ca^2+^ from vacuoles and other organelles to the cytosol (Pei et al., 2000; Wang et al., 2013; Wu et al., 2020).

ROS production via RBOHs is regulated by a suite of different kinases at the whole leaf, tissue, cellular and organellar levels. The Ca^2+^ sensor CALCINEURIN B-LIKE proteins (CBL1; CBL9) interact with CBL-INTERACTING PROTEIN KINASE (CIPK11; CIPK26) to activate RBOHF (Drerup et al., 2013). In contrast, RBOHD is the primary apoplastic ROS generator in response to biotic stimuli and mechanical wounding (Morales et al., 2016; Torres et al., 2002). Regulators of RBOHD activity include several immunity-associated protein kinases (Lee et al., 2020; Li et al., 2014; Zhang et al., 2018), and calcium protein kinases (CPKs). CPK1, CPK2, CPK4, CPK5, CPK6 and CPK11 can activate both RBOHD and RBOHF in an immunocomplex kinase assay (Gao et al., 2013), and at least for CPK5, a direct interaction and activation of RBOHD has been confirmed (Dubiella et al., 2013). In guard cells, OST1 activating RBOHF is thought to be the primary pathway for apoplastic ROS production (Sirichandra et al., 2009; Wu et al., 2020). However, the links between ROS production and CPKs as well as the interaction with retrograde signals and the production of apoplastic ROS in guard cells remain elusive, if not illogical from a spatial context.

The aforementioned leaf chloroplast PSI ROS can trigger other chloroplast-to-nucleus retrograde signaling pathways such as methylerythritol cyclodiphosphate (MEcPP) and SAL1-PAP (3’phosphoadenosine 5’phosphate), the latter of which functions in drought and high light stress (Gonzalo M Estavillo et al., 2011; Xiao et al., 2012). These abiotic stresses promote chloroplast PSI ROS and redox poise which inhibit plastidial SAL1 activity (Chan et al., 2016a), thus allowing accumulation of its catabolic substrate PAP in the cell. In the nucleus, PAP inhibits 5’ to 3’ exoribonucleases to alter RNA polymerase II function. This leads to read through accumulation of downstream genes in tandem repeats, resulting in accumulation of stress-responsive transcripts including H_2_O_2_ scavengers (Crisp et al., 2018, 2017). Consequently, *sal1* mutants with constitutively over-accumulated PAP show lower foliar ROS levels, particularly in the vascular bundle (Estavillo et al., 2011).

In guard cells, the SAL1-PAP pathway functions in parallel to the canonical ABA signaling (RCAR/PYL-OST1-PP2C) pathway to induce stomatal closure, at least in part via transcriptional upregulation of CPKs which converge upon, and activate, OST1 target proteins such as the SLAC1 anion channel (Pornsiriwong et al., 2017). Intriguingly, PAP appears to have contrasting effects on ROS in different cell types. While constitutive PAP accumulation leads to decreased ROS in vascular bundles (Estavillo et al., 2011), exogenous PAP application induces ROS in guard cells (Pornsiriwong et al., 2017). Constitutively accumulated PAP also rescues ABA-induced ROS burst in *ost1 sal1* guard cells (Pornsiriwong et al., 2017). However, the mechanism was not investigated, beyond establishing that most downstream canonical ABA-regulated processes, such as ion fluxes, were activated by PAP in *ost1* mutants.

These contrasting roles for PAP as an initiator and repressor of ROS signaling in different cell types raise the hypothesis tested herein that chloroplasts and their retrograde signals can function differently depending on cellular context. Further, despite the parallels in chloroplast-sourced retrograde ROS signaling and apoplast-sourced secondary messenger ROS signaling, there is a distinct lack of knowledge how these two processes are intertwined in stress signaling. In this work we explore how SAL1-PAP retrograde signaling intersects with whole-leaf ROS responses such as basal plasmodesmal permeability, flg22-induced ROS responses and tolerance to ozone-derived apoplastic ROS. To define how PAP, and consequently chloroplastic signals, can intersect with ROS signaling, we use the guard cell as a model to study the specializations of each pathway. This includes considerations of the sites of ROS accumulation, phosphorylation targets of CPKs and the extent to which ABA-mediated and PAP-mediated signaling require chloroplast functionality as well as retrograde signaling for stomatal closure.

## Results

### Subcellular guard cell ROS compartmentalization is mediated by retrograde signaling for stomatal closure

Previously, we showed that ROS signaling was restored by PAP in ABA signaling mutants, but we did not determine the subcellular source of ROS nor its regulation (Pornsiriwong et al., 2017). Herein we examined the spatial distribution of ROS in response to exogenous PAP application using H_2_DCFDA which acts as a general ROS marker by reacting with H_2_O_2_ and other radicals such as OHꞏ (Akter et al., 2021). H_2_DCFDA has been used to visualize ROS accumulation in whole leaves and subcellular compartments of guard cells (Postiglione and Muday, 2023; Watkins et al., 2017; Zandalinas et al., 2020).

We observed that PAP and ABA elevated ROS in multiple guard cell subcellular compartments, with a notable and significant increase in chloroplasts (Fig. 1A, B). ABA treatments induced chloroplastic ROS to a lower extent than PAP (Fig. 1B). The observed increase in H_2_DCFDA fluorescence by approximately 1.2 - 1.5-fold in response to 10 min treatment with PAP or ABA (Fig. 1A, B) is similar to that previously reported for H_2_DCFDA fluorescence in *Arabidopsis thaliana* (*Arabidopsis*) and tomato guard cells after 15 min of ABA treatment (Postiglione and Muday, 2023; Watkins et al., 2017). We also visualized ROS accumulation in PAP-treated guard cells using the H_2_O_2_-specific fluorescent dye Peroxy Orange 1 (PO1) (Dickinson et al., 2010) and found comparable PAP-induced fluorescence increases in the guard cell between H_2_DCFDA and PO1 (Fig. 1A, Fig. S1), consistent with previous observations (Postiglione and Muday, 2023; Watkins et al., 2017; Zandalinas et al., 2020). Consequently, we conclude that PAP induces accumulation of ROS, including H_2_O_2_, in guard cells.

**Figure 1.**
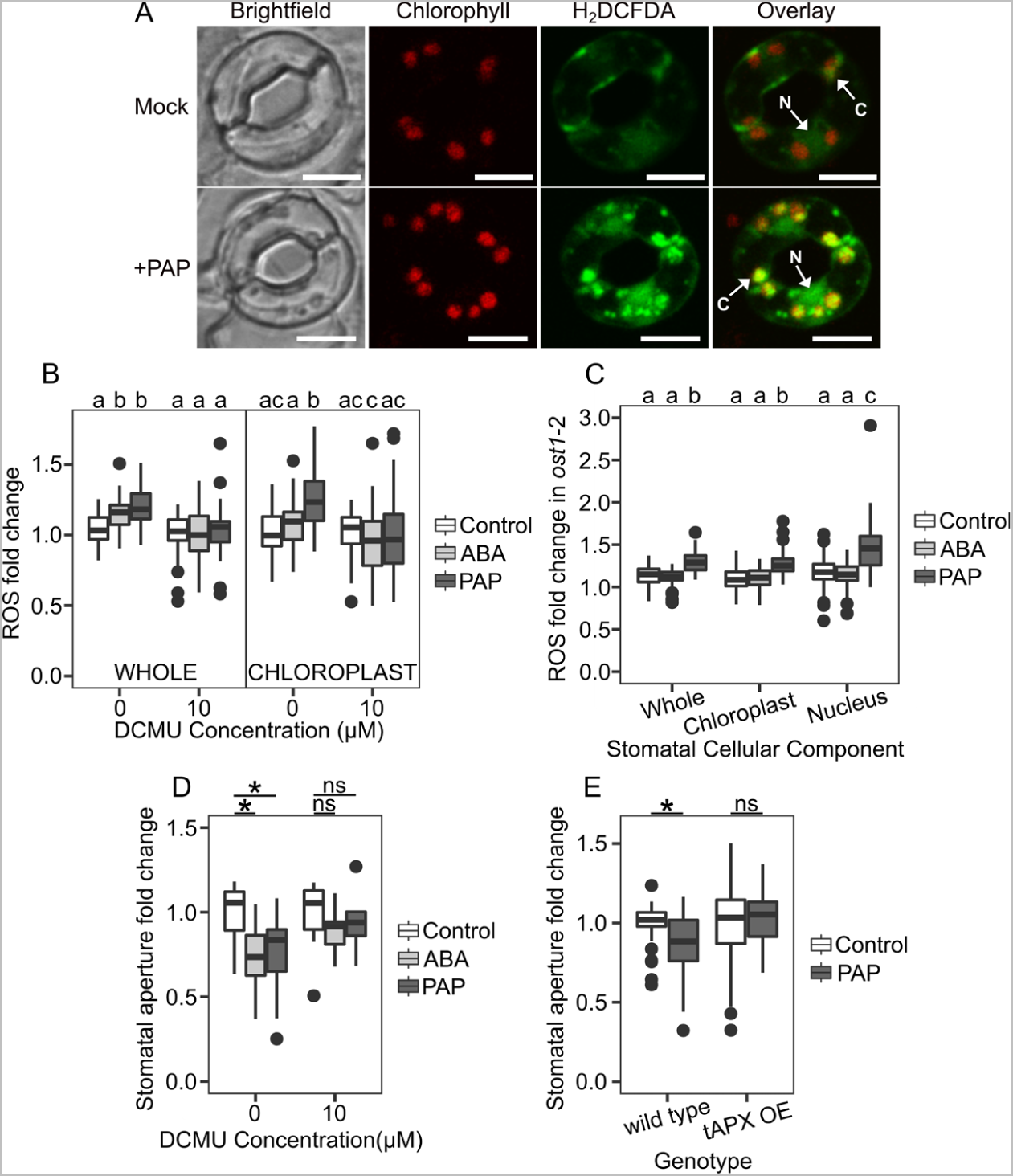
Subcellular compartmentalization of ROS intersects with retrograde PAP signaling. (A) Localization of ROS within guard cells treated with mock or PAP, with arrows indicating ROS at chloroplasts (C) and the nucleus (N). Scale bars = 10 µM. (B) and (E) show relative change in fluorescence (compared to control) in different parts of the stomata after treatment (100 µM ABA or 100 µM PAP). Fluorescence is an indication of ROS levels as per the ROS dye H_2_DCFDA. (B) shows the relative change in fluorescence in whole guard cells or guard cell chloroplasts after treatments with or without 10 µM DCMU. Each condition contains a minimum of three wild type plants with n ≥ 49 stomata per treatment combination. For ‘Whole’, factors ‘DCMU’ and ‘Treatment’ significantly impacted fluorescence with a significant interaction between the two (ANOVA; F =59.6, df=1, p<0.001; F=12.0, df=2, p<0.001; F = 7.3, df=2, p<0.001). Significant differences from the respective control denoted by a and b. For ‘Chloroplast’, factors ‘DCMU’ and ‘Treatment’ significantly impacted fluorescence, with a significant interaction between the two (ANOVA; F =32.7, df=1, p<0.001; F=23.7, df=2, p<0.001; F=11.6, df=2, p<0.001). Significant differences from the respective control denoted by a, b or c. For (C), each treatment/subcellular component combination contains a three *ost1*-2 plants with a minimum of n ≥ 40 stomata. ‘Treatment’ and ‘Part’ significantly impacted fluorescence (ANOVA; F=68.8, df=2, p<0.001 and F=13.4, df=2 p<0.001), with a significant interaction between the two (ANOVA; F=3.2, df=4, p<0.05). Significant differences denoted by a, b and c. (D) Stomatal aperture in response to 100 µM ABA or 100 µM PAP with 0 or 10 µM DCMU, fold change relative to DCMU control (i.e., 0 DCMU or 10 µM DCMU). Each condition contains a minimum of three wild type plants with a minimum of n ≥ 27 stomata per treatment combination. ‘DCMU’ and ‘Treatment’ significantly impacted closure (ANOVA; F=7.7, df=1, p<0.01; F=4.2, df=2, p<0.05), with significant differences from respective control determined denoted by *, p < 0.05. (E) Stomatal closure in response to 100 µM PAP in tAPX OE. Each condition contains a minimum of three biological replicates with a minimum of n ≥ 41 stomata per treatment combination. ‘Genotype’ significantly impacted closure (ANOVA; F= 5.4, df=1, p<0.05), with a significant interaction between ‘Genotype’ and ‘Treatment’ (ANOVA; F=4.9, df=1, p<0.05). Significant differences from the genotype control denoted by *, p<0.05.

To test whether the effect of PAP on chloroplast ROS requires ABA signaling components, we investigated PAP treatment in an *ost1-2* background which is deficient in ABA signaling and consequently ABA-induced stomatal closure (Mustilli et al., 2002; Pornsiriwong et al., 2017). PAP still induced chloroplast ROS in *ost1-2* (Fig. 1C) while ABA did not, suggesting that PAP acts independently of ABA in the induction of chloroplast ROS.

Given PAP is reported to lower, not raise, chloroplast ROS in mesophyll and bundle sheath leaf tissues we investigated further the interaction between PAP and chloroplastic ROS using biochemical and genetic tools. The chloroplast photosynthetic electron transport chain is a major source of ROS in chloroplasts (Waszczak et al., 2018). Therefore, we attempted to attenuate chloroplastic ROS production using different concentrations of the photosynthesis inhibitor 3-(3-4,-dichlorophenyl)-1,1-dimethylurea (DCMU) on *Arabidopsis* epidermal peels. As expected, we observed incremental reduction of the effective PSII quantum yield (Y(II)) of mesophyll tissue in response to the increasing concentrations of DCMU (Fig. S2); whereas Y(II) in the guard cells was below the detection threshold. We used this decrease of Y(II) in the mesophyll to infer that Y(II) was similarly decreased in the chloroplasts of guard cells given the similarity of chloroplast photosynthesis in both cell types (Lawson et al., 2003, 2008). Inhibition of guard cell photosynthesis and chloroplastic ROS production at 10 µM DCMU decreased both ROS accumulation in guard cells and guard cell chloroplasts treated with ABA or PAP (Fig. 1B). This decreased ROS correlated with impaired ABA- or PAP-induced stomatal closure when co-treated with a range of DCMU concentrations (Fig. 1D, S3). Intriguingly, this PAP-induced increase in ROS level driven by the light reactions of photosynthesis was not associated with any changes to the low light regime; that is, there was no requirement for co-treatment with high light as would be typically expected for chloroplastic ROS production.

To further test if chloroplastic PSI ROS contributes to PAP-mediated stomatal closure, we utilized a thylakoid ascorbate peroxidase (tAPX) over-expressor (tAPX-OE) line (Murgia et al., 2004). This transgenic line has high levels of tAPX activity for catalyzing the reduction of chloroplastic H_2_O_2_ to water. We hypothesized that increased scavenging of chloroplastic PSI ROS would prevent PAP-mediated stomatal closure. Indeed, PAP-mediated stomatal closure was inhibited in tAPX-OE in comparison to wild type (Fig. 1E).

As chloroplast-nuclear proximity and H_2_O_2_ transfer between these compartments in photosynthetic tissues have been proposed as a key signaling mechanism (Exposito-Rodriguez et al., 2017), we examined these processes in the context of PAP signaling in guard cells. We first verified that PAP also induces an increase in ROS levels in the nucleus of guard cells by overlay of H_2_DCFDA fluorescence with that of the nuclear marker DAPI (Fig. S4). Given the finding that PAP can induce chloroplastic ROS in *ost1*-2 (Fig. 1C), we quantified H_2_DCFDA fluorescence in both chloroplasts and the nucleus in parallel and found that PAP treatment increases ROS levels in the nucleus to a higher degree than in the chloroplasts (Fig. 1C).

We then examined whether there was a spatial aspect to this chloroplast-nuclear ROS increase. Specifically, did chloroplasts associated with the nucleus respond differently to PAP compared to chloroplasts elsewhere in the cell? First, we determined whether PAP treatment altered the localization of chloroplasts within a guard cell. Of the three to six chloroplasts per single guard cell, one or two chloroplasts were typically associated with the junction between two guard cells, two to three chloroplasts with the nucleus, and one or none in neither location (classed as ‘other’, Fig. S5). The number of chloroplasts associated at the junction, nucleus or ‘other’ location was not affected by PAP treatment (Fig. S5). Significantly, PAP induced ROS to a similar extent in all guard cell chloroplasts regardless of location, including those associated with the nucleus (Fig. S6). Collectively, the pattern of PAP-induced ROS accumulation in both chloroplasts and the nucleus of guard cells is very similar to that induced by ABA (Postiglione and Muday, 2023; Watkins et al., 2017). Therefore, our results indicate that like ABA, PAP induces ROS production in guard cell chloroplasts, with the ROS capable of accumulating to high levels in the nucleus.

### Apoplastic ROS production by RBOHD is required for PAP-mediated signaling processes

While apoplastic ROS is required for stomatal closure mediated by ABA (Rodrigues et al., 2017), the role and genetic components for this process have not been identified for PAP-mediated stomatal closure. We found that the double mutant *rbohDrbohF* (*rbohDF*) did not close stomata in response to PAP and had the expected impaired ABA-mediated stomatal closure (June M. Kwak et al., 2003) (Fig. 2A). Interestingly, we found that the single knockout *rbohF* mutant was responsive to PAP but had completely impaired ABA-mediated stomatal closure (i.e., no different to the control). In contrast, ABA induced stomatal closure in *rbohD*, but PAP could not. Indeed, while ABA induced a significant increase in ROS of *rbohD* guard cells, PAP treatment on *rbohD* did not (Fig. 2B); this was also the case for when chloroplastic ROS was measured by quantifying H_2_DCFDA fluorescence specifically in areas overlapping with chlorophyll fluorescence (Fig. 2B). This indicates that ABA induces ROS via RBOHF whereas PAP acts through RBOHD, which marks an important delineation between ABA- and PAP-mediated stomatal closure.

**Figure 2.**
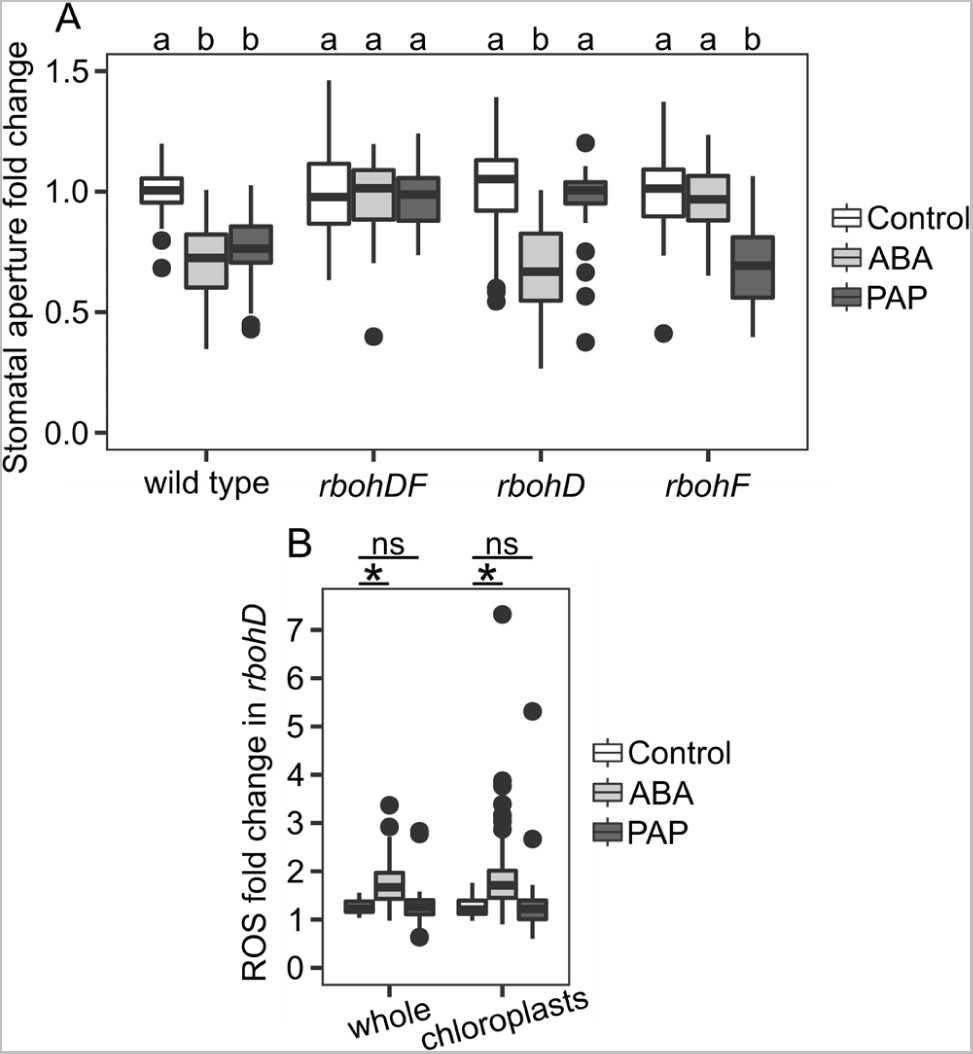
*rboh* mutants are differentially perturbed in ABA and PAP-mediated stomatal closure. (A) Stomatal closure of *rboh* mutants in response to 100 µM ABA or 100 µM PAP. Each condition contains a minimum of three biological replicates with a minimum of n ≥ 34 stomata per treatment combination. ‘Genotype’ and ‘Treatment’ significantly impacted closure with an interaction between the two (ANOVA; F=15.8, df=3, p<0.001; F=38.6, df=2, p<0.001; F=18.1, df=6, p<0.001). Significant differences to the respective genotype control denoted by a or b. (B) Relative change in fluorescence in whole or chloroplasts of *rbohD* guard cells as an indication of ROS levels. Each condition contains a minimum of three biological replicates with a minimum of n ≥ 38 stomata per treatment combination. ‘Treatment’ significantly impacted fluorescence in both ‘whole’ and ‘chloroplasts’ datasets (ANOVA; F=16.5, df=2, p<0.01; F=7.4, df=2, p<0.05), with significant differences from the controls denoted by *, p<0.05.

The requirement of RBOHD for PAP-induced chloroplastic ROS (Fig. 2B) would suggest that production of plastidial ROS is initiated by extracellular ROS in this signaling cascade. We then tested the reverse hypothesis, that is, apoplastic ROS production requires chloroplastic ROS. We utilized a modified form of H_2_DCFDA, Oxyburst Green H2HFF-BSA, which is restricted to the apoplast and thus acts as a general ROS marker specifically in this compartment (Miller et al., 2010). PAP treatment induced an increase in apoplastic Oxyburst Green H2HFF-BSA fluorescence in guard cells compared to mock treatment as expected (Fig. S7). Inhibition of chloroplast ROS production led to a visible decrease in apoplastic Oxyburst Green H2HFF-BSA fluorescence in guard cells co-treated with PAP and DCMU (Fig. S7). Taken together, our data suggest that the effect of PAP on subcellular ROS is multifaceted, with PAP-induced ROS production in chloroplasts and the apoplast influencing each other.

### PAP-induced CPKs activate RBOHD and SLAC1 thereby simultaneously impacting ROS and anion currents in stomatal closure

The induction of RBOHD activity for ROS production is thought to occur both by calcium transients and protein kinase activation (Lee et al., 2020; Li et al., 2014; Zhang et al., 2018). Intracellular calcium binds to the EF-hand motifs of RBOH, and calcium influx can lead to the activation of several kinases such as the CPKs, which can phosphorylate RBOHD at the N terminus (Dubiella et al., 2013). Considering PAP requires RBOHD to initiate stomatal closure, we investigated potential CPK candidates shown to be transcriptionally up-regulated by PAP (Pornsiriwong et al., 2017) as enhancers of RBOHD-mediated ROS production.

To determine whether these candidate CPKs could enhance the ROS-producing activity of RBOHD, we used heterologous expression in human embryonic kidney 293T (HEK293T) cells. These cells lack endogenous ROS production activity under experimental conditions employed herein (Bánfi et al., 2001) and are routinely used to investigate post-translational regulation of RBOH activity (Chu et al., 2023; Kaya et al., 2014). Measurement of ROS in these cells by luminol-amplified chemiluminescence showed that the single transfections of CPKs alone did not differ significantly to the non-transfected control, indicating that the CPKs alone could not induce ROS production (Fig. S8). HEK293T cells were then co-transfected with *3FLAG-RBOHD* and either *CPK-3Myc*, or *3Myc-GFP* as a negative control (Fig. 3A). RBOHD protein levels were homogenous between co-transfections, indicating that any differences in ROS production would be due to an additional protein partner (Fig. S9). Upon stimulation with ionomycin which supplies Ca^2+^ as a co-factor of RBOH proteins, the negative control HEK293T cells co-expressing *3FLAG-RBOHD* and *3Myc-GFP* exhibited only a small increase in luminescence, indicating minimal ROS production. By contrast, HEK293T cells co-transfected with 3*FLAG-RBOHD* and one of either CPK12, CPK28, CPK32 or CPK34 showed significantly enhanced RBOHD-mediated ROS production compared to the negative control (Fig. 3A). CPK13 did not induce significantly different ROS production compared to the control, despite CPK13 being expressed to higher levels than CPK28 or CPK32 in the HEK293T cells (Fig. S9). Overall, these results demonstrate that multiple, yet specific, CPKs that are transcriptionally up-regulated by PAP can activate RBOHD for PAP-mediated ROS functions.

**Figure 3.**
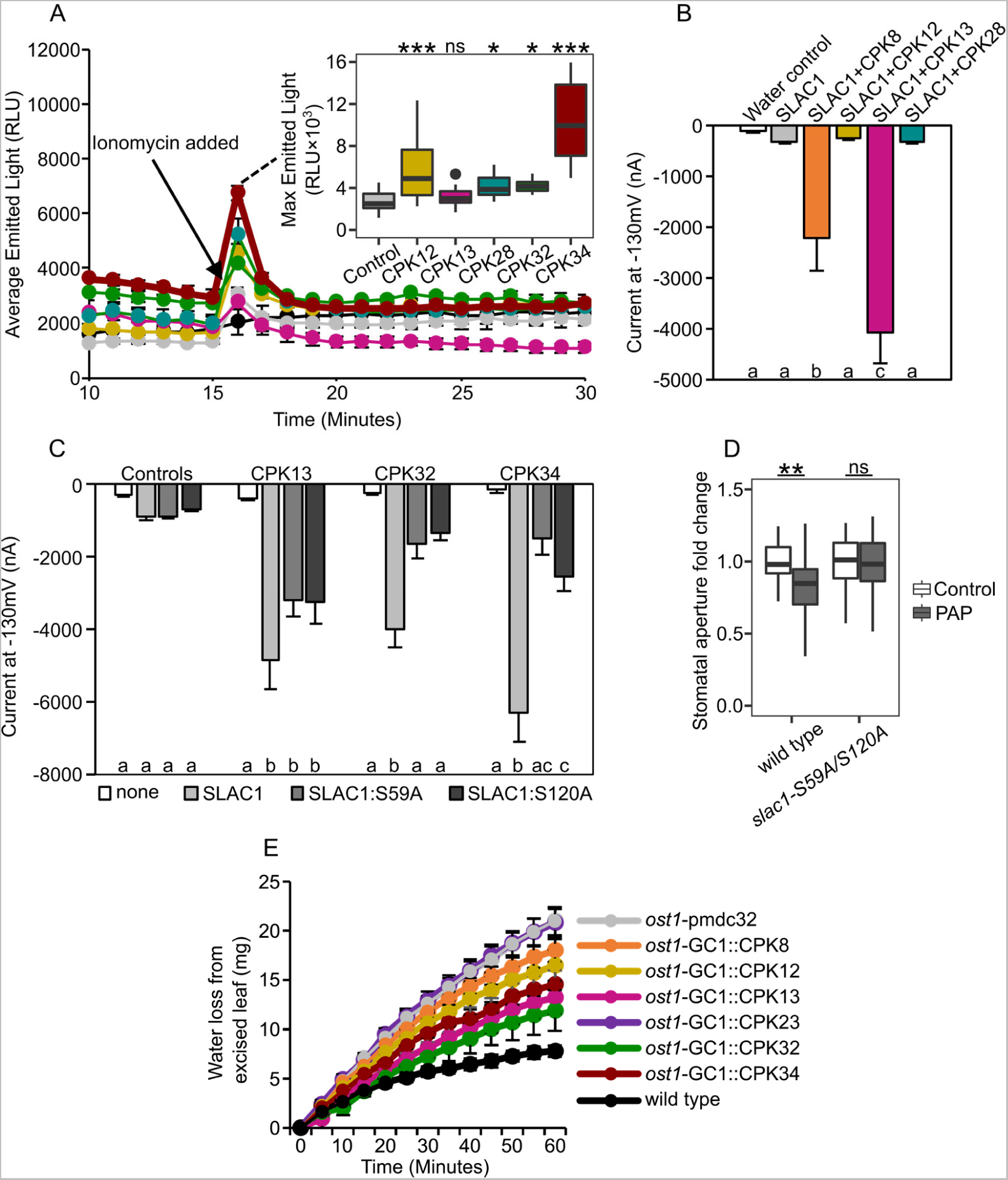
Different PAP-induced CPKs enhance RBOHD-mediated ROS production and SLAC1 anion currents. (A) ROS production of HEK293T cells transiently expressing CPKs with RBOHD. After 15 min 1 µM ionomycin was added to the medium. Values represent mean ± SEM for either three non-transfected replicates (shown in black), RBOHD co-transfected with GFP (control) or designated CPK. The experiment was repeated in at least four independent experiments with similar results; boxplot shows average max emitted light with data pooled from separate experiments. Control n = 24, CPK12 n = 18, CPK13, 28, 32, and 34 n = 12. Factor ‘Gene’ expressed with RBOHD significantly impacted the max emitted light after ionomycin was added (ANOVA; F=32.3, df=5, p<0.001), with significant differences from the control denoted as * for p<0.05 or *** for p<0.0001. (B) CPK8, CPK12, CPK13 and CPK28 activation of SLAC1 anion channel in oocytes. Values are means of four to eight oocytes ± SEM. Anion channel activity shown as steady state currents activated at -130 mV, with significant differences between injected combinations (ANOVA; F=3.5, df=5, p<0.01), denoted by a, b and c. Experiment was repeated on positive candidates in two other batches of oocytes. See Fig. S7 for full trace. (C) Anion currents of mutated phosphosites in SLAC1 when co-expressed with CPK13, CPK32 and CPK34 in oocytes. Values are from a minimum of five oocytes per combination ± SEM. Factors ‘Type of SLAC1 expressed’ and ‘CPK’ expressed significantly impacted the anion currents, with a significant interaction between the two (ANOVA; F=70.8, df=3, p<0.001; F=33.5, df=3, p<0.001; F=16.8, df=9, p<0.001). Significant differences with a given CPK expressed denoted by a, b and c. (D) Stomatal aperture of *slac1-S59A/S120A* mutant in response to 100 µM PAP. Each has a minimum of three biological replicates with a minimum of n ≥ 31 stomata per genotype/treatment combination. Factors ‘Genotype’ and ‘Treatment’ significantly impacted closure with a significant interaction between the two (ANOVA; F=6.3, df=1, p<0.05; F=6.4, df=1, p<0.05; F = 5.5, df=1, p<0.05). Significant differences from the genotype control denoted by **, p<0.01. (E) Water loss assay in excised leaves of *ost1* complemented with guard-cell promoter expressed CPKs. Independent transgenics lines, where n = 8, 17, 14, 9, 13, 6, 14 and 11 respectively for *ost1*-pmdc32, *ost1*-GC::CPK8, *ost1*-GC::CPK12, *ost1*-GC::CPK13, *ost1*-GC::CPK23, *ost1*-GC::CPK32, *ost1*-GC::CPK34, and wild type.

Interestingly, the RBOHD-activating kinases include CPK32 and CPK34, which we have previously identified as CPKs capable of activating SLAC1 in *Xenopus laevis* oocytes (Pornsiriwong et al., 2017). This raises the question of which, amongst the repertoire of PAP-upregulated CPKs, might target RBOHD and SLAC1 together or individually. We therefore tested CPK8 [another CPK transcriptionally upregulated in response to PAP and known to have roles in stomatal closure (Zou et al., 2015)], CPK12, CPK13 and CPK28 for SLAC1 activation using the *Xenopus* heterologous expression system (Fig. 3B, S10-12). CPK12 could not activate SLAC1 in oocytes, consistent with findings by Brandt et al. (2012), nor could CPK28 elicit SLAC1 anion currents. However, both CPK8 and CPK13 could elicit SLAC1 anion currents, albeit to varying degrees in four independent experiments. Overall, different members of the PAP-induced CPKs target RBOHD and SLAC1 either individually or collectively (Table S1).

Different kinases phosphorylate different serine sites on SLAC1 (Brandt et al., 2015, 2012; Geiger et al., 2009). Serine 120 (S120) is phosphorylated by OST1, while CPK6 phosphorylates S59 (Brandt et al., 2015). To define whether some of our identified CPK candidates target these phosphorylation sites, S59 and S120 of SLAC1 were individually or both mutated to alanine. SLAC1:S59A and SLAC1:S120A were individually co-expressed in oocytes with CPK13, CPK32 and CPK34 (Fig. 3C, S12). Contrary to the published literature on OST1 and CPKs phosphorylating distinct sites on SLAC1, either one of the S59A or S120A mutations in SLAC1 on its own led to a reduction in anion currents for CPK32 and CPK34. When we mutated both serine sites (i.e., SLAC1-S59A:S120A), this led to a further reduction in anion currents for CPK13 (Fig. S12). We validated these findings *in planta* by testing whether different *Arabidopsis* SLAC1 mutants responded to PAP. As expected, PAP and ABA did not close stomata of *slac1-4* lacking the SLAC1 protein (Fig. S13). Importantly, the *slac1-S59A/S120A* double mutant which expresses SLAC1 containing mutated serine sites, did not close stomata when treated with PAP (Fig. 3D).

Considering these PAP-related CPKs could activate both RBOHD and the anion channel SLAC1 individually, or both, thereby simultaneously impacting ROS and anion currents in stomatal closure, we wanted to determine whether they could compensate for the lack of OST1, an important kinase that activates SLAC1 and RBOHF (Acharya et al., 2013; Sirichandra et al., 2009). We transgenically over-expressed the different CPKs in the *ost1* background and performed a water-loss assay on excised individual leaves of these transgenic lines. We included CPK23, which is a known activator of SLAC1 involved in the ABA signaling pathway (Geiger et al., 2010) but not transcriptionally PAP-induced; expressing CPK23 in *ost1* did not alter the water-loss rate when compared to the empty-vector *ost1* control (Fig. 3E). In contrast, the overexpression of PAP-induced CPKs in *ost1* could reduce the water loss in comparison to empty-vector *ost1* to varying degrees (Fig. 3E). This indicates that PAP-induced CPKs, in particular CPK32 and CPK34, can, when expressed singularly, partially compensate for the lack of OST1 *in vivo* by modulating RBOHD and/or SLAC1 activity (Table S1).

### PAP intersects with ROS signaling across diverse cell types and processes

ROS, in particular H_2_O_2_, are critical secondary messengers in many cell types and cellular processes. Therefore, we explored the extent to which PAP might intersect with ROS beyond the guard cell. First, we investigated whether the characteristic ROS production induced by the pathogen elicitor flg22 (Li et al., 2014; Morales et al., 2016; Torres et al., 2002) is altered in *sal1* leaves as a whole, given that both flg22 and now PAP (Fig. 2A) have been shown to act via RBOHD in guard cells. Flg22 induced a much more pronounced ‘initial’ ROS burst in *sal1* in comparison to wild type (Fig. 4A), while the negative control *rbohD* did not produce any ROS.

**Figure 4.**
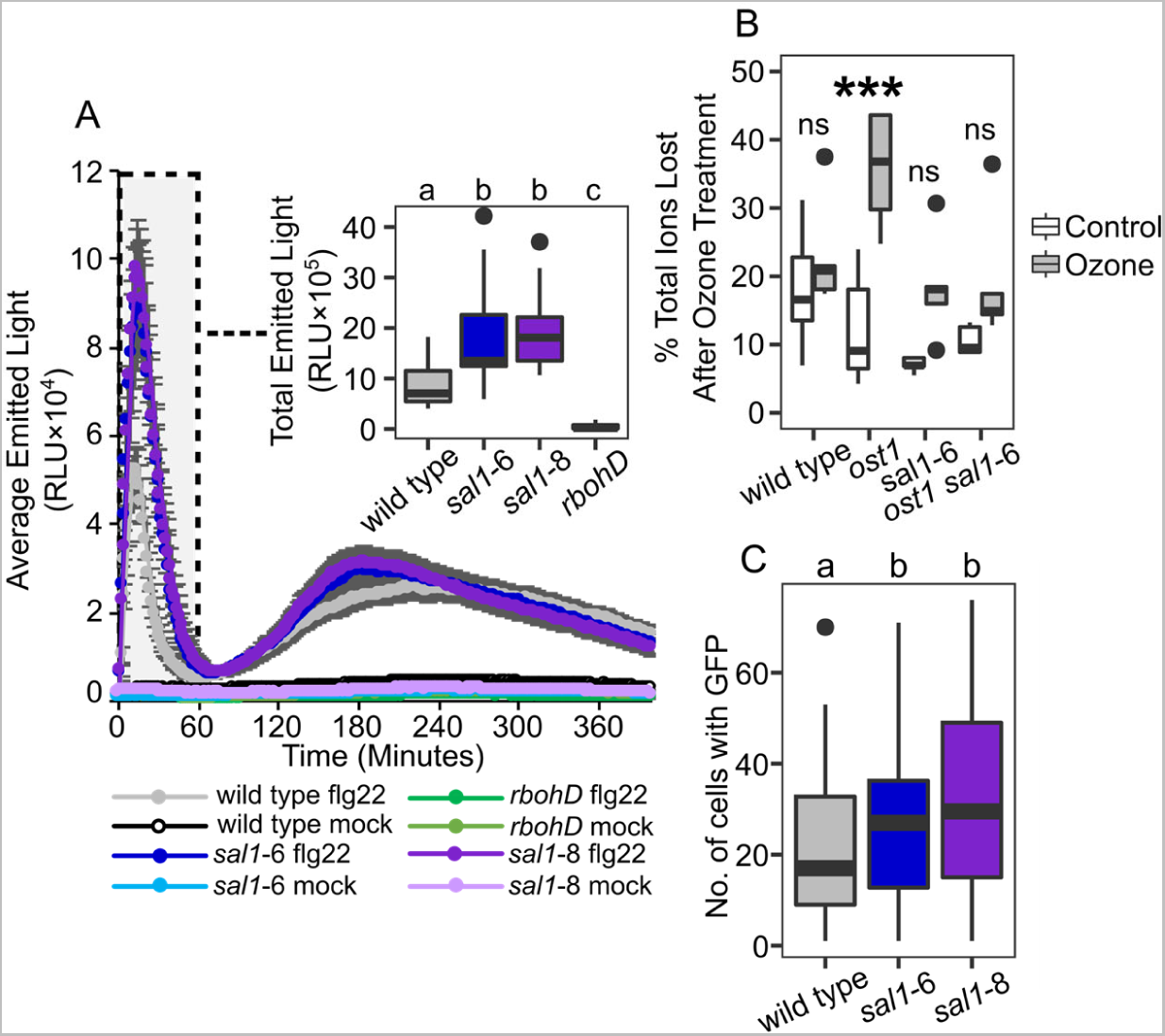
Altered cellular ROS processes in *sal1 mutant.* (A) Flg22 chemiluminescence response kinetics in Arabidopsis *sal1* mutant leaf discs. Average of five-week-old plants ± SEM (eight per genotype and treatment), with data combined from two separate experiments (n=16). Boxplot shows total emitted light over 60 minutes for wild type, *sal1*-6 and *sal1*-8 and *rbohD* in response to flg22. Independent factor ‘Genotype’ was determined as significant (ANOVA; F=36.6, df=3, p<0.001); significant differences between genotypes denoted by a, b and c. (B) Electrolyte leakage measurements from plants harvested after six-hour exposure to ozone, 24 hours after start of ozone exposure. Values are plotted as a % total ion lost, with n = 5 per genotype and treatment. Factors ‘Genotype’ and ‘Treatment’ had a significant impact on total ion lost (ANOVA; F=4.7, df=3, p<0.01 and F=25.5, df=1, p<0.001), with significant differences between treatments within a genotype denoted by ***, p<0.001. (C) Microprojectile bombardment into leaf tissue of *sal1*-6, *sal1*-8 and wild type show *sal1* mutants have more GFP movement to neighboring cells than wild type. Number of bombardment sites counted ≥84 per genotype, three leaves × three different plants per genotype. Experiment repeated with similar significant trends found. Significance differences between genotypes indicated by a and b.

Ozone enters leaves via stomata, and is readily converted to secondary ROS including H_2_O_2_, super oxide and hydroxyl radical in the apoplast (Vainonen and Kangasjärvi, 2015). The whole-leaf response to ozone requires functional OST1 (Vahisalu et al., 2010) and perturbation of this response can result in excessive damage to vascular tissue (Vahisalu et al., 2008). Considering our previous finding that constitutive PAP signaling via *sal1* restores stomatal closure in *ost1* (Pornsiriwong et al., 2017), we investigated whether *ost1 sal1*-6 has restored ozone tolerance. The ozone-tolerant wild type (ecotype Col-0) and ozone-sensitive *ost1* plants showed the expected phenotypes, with lesions formed in the midrib of all leaves of *ost1* (Fig. S14). Interestingly, fewer lesions developed on *ost1 sal1*-6; and said lesions were primarily contained to the older leaves in the lower part of the rosette. In accordance with this, whole plant electrolyte measurements showed that the significantly greater extent of ozone-induced ion leakage in *ost1* was decreased to wild type levels in *ost1 sal1-6* (Fig. 4B).

To identify putative mechanisms enabling this rescued ozone tolerance, we performed Gene Ontology (GO) enrichment analysis (Tian et al., 2020) on our previously published transcriptome of *ost1 sal1-6* (Pornsiriwong et al., 2017). We found significant enrichment in differentially expressed genes relating to oxidation-reduction and phosphorylation, which include ROS homeostasis enzymes (e.g. ascorbate peroxidases, glutathione peroxidases) and signaling kinases (e.g. MAP kinases, CPKs) respectively (Table S2). Many of these genes are established components of the plant cellular response to ROS (Waszczak et al., 2018). Therefore, collectively our data indicate that PAP accumulation restores ozone tolerance, and thereby ROS responses, in whole leaves lacking OST1.

We finally examined the effect of constitutive PAP accumulation on another cellular process which utilize ROS production and/or signaling; namely cell-to-cell permeability via plasmodesmata. The aperture of plasmodesmata can be dynamically regulated by ROS in roots treated with exogenous H_2_O_2_ (Rutschow et al., 2011), and plasmodesmata have been implicated in systemic leaf-to-leaf ROS signaling (Fichman et al., 2021). The *sal1* mutant alleles *sal1-*6 and *sal1-*8 showed more open plasmodesmata in comparison to wild type, as indicated by greater diffusion of green fluorescent protein (GFP) from the initial particle bombardment site into neighboring cells (Fig. 4C). Interestingly we identified several genes related to plasmodesmata aperture regulation that were also differentially expressed in *sal1*-8 compared to wild type (Table S3). Collectively, these results for stomata, flg22, ozone and plasmodesmata suggest the potential for PAP to intersect with ROS signaling and homeostasis in diverse cellular processes.

## Discussion

Unanswered questions in the fields of ROS and retrograde signaling revolve around functional redundancy versus specificity of signals, as well as a less commonly considered quandary which is the spatio-temporal intersection and neofunctionalization of the different signals in specialized cells. For instance, if chloroplastic H_2_O_2_ itself is exported to the nucleus for signaling roles (Exposito-Rodriguez et al., 2017), why do plant cells require other H_2_O_2_-triggered retrograde signals? Furthermore, signaling roles for chloroplastic and apoplastic ROS have conventionally been considered separately; thus, whether and how they intersect has remained unclear. A more fundamental question is if retrograde signaling is an important cellular mechanism from an evolutionary perspective, how does it intersect with other stress signaling pathways? Does this differ in specialized cells, such as guard cells versus established roles in mesophyll cells or vasculature bundles, as is the case for hormones and secondary messages? Collectively, our results suggest the potential for PAP to intersect with ROS signaling and homeostasis in diverse cellular processes.

### PAP induces ROS in multiple subcellular compartments, including chloroplasts

We observed that both PAP and ABA induced ROS in chloroplasts, nuclei and the apoplast of the guard cells, with similar patterns of subcellular fluorescence being observed for both the general ROS marker H_2_DCFDA and the more H_2_O_2_-specific PO1 (Fig. 1, Fig. S1). Our results are consistent with recent reports of ABA inducing ROS accumulation in chloroplasts, nuclei and mitochondria of *Arabidopsis* and tomato guard cells (Postiglione and Muday, 2023; Watkins et al., 2017). The authors also found similar extents of ROS-responsive fluorescence changes between H_2_DCFDA and PO1, as well as with the genetically-encoded H_2_O_2_ biosensor protein roGFP2-Orp1 (Postiglione and Muday, 2023).

Our observed increases in H_2_DCFDA fluorescence by approximately 1.5-fold in chloroplasts and nuclei of wild type guard cells after 10 min treatment with PAP or ABA, when juxtaposed against the unchanged fluorescence in ABA-treated *ost1* (Fig. 1A, B) indicate that these localized ROS increases are genuine effects of signaling cascades in the guard cells (Fig. 1A, B) (Postiglione and Muday, 2023; Watkins et al., 2017). This suggests that increases in chloroplast and nuclear ROS are utilized across different signaling cascades for stomatal regulation (Postiglione and Muday, 2023; Watkins et al., 2017).

The inhibition of PAP-induced stomatal closure by DCMU or tAPX over-expression (Fig. 1B, D, E, S3) identifies PSI and the requirement for functioning photosynthetic electron transport chain as likely origins of the chloroplastic ROS required for stomatal closure. It seems that chloroplastic PSI ROS and photosynthesis *per se* play important roles in guard cell signaling, with PAP, ABA and apoplastic ROS among the triggers for increases in photosynthetic-generated ROS [Figure 1 and (Postiglione and Muday, 2023; Watkins et al., 2017)]. Presumably there is a synergistic interaction between chloroplastic ROS and RBOH-mediated ROS for stomatal closure, such as facilitating calcium release thereby promoting activation of ion channels via CPKs (Demidchik and Shabala, 2018; Sierla et al., 2016). Consistent with this assumption, suppression of chloroplast ROS with DCMU visibly decreased PAP-induced apoplastic ROS (Fig. S7).

Of note, we also observed a concurrent increase in nuclear ROS in response to PAP (Fig. 1C), which is consistent with an emerging understanding of a role for transcription in stomatal function (Pornsiriwong et al., 2017; Simeoni et al., 2022). Guard cells also have a higher proportion of chloroplasts associated with the nucleus (Fig. S5) compared to mesophyll cells, where direct H_2_O_2_ transfer from chloroplasts to the nucleus was reported (Exposito-Rodriguez et al., 2017); however we saw no evidence for PAP altering the interactions of plastids with nuclei. While well beyond the scope of this study one can speculate that chloroplast-sourced ROS oxidizing guard cell nuclear proteins may alter protein interactions, trafficking, conformation and function (Couturier et al., 2013; Wei et al., 2020). Mining of recent proteomics studies identifying proteins with H_2_O_2_-sensitive cysteine residues (Huang et al., 2019; Wei et al., 2020) revealed several nuclear proteins involved in stomatal regulation (Chen et al., 2021; Tõldsepp et al., 2018). This list includes CO_2_ INSENSITIVE 1 (CIS1) which is required for elevated intracellular bicarbonate-induced activation of S-type anion channel currents for CO_2_-induced stomatal closure (He et al., 2018). The diversity of these proteins opens new research avenues for understanding chloroplastic ROS signaling.

### How do ROS from different subcellular compartments intersect for stomatal closure?

Apart from direct ROS/retrograde signaling targets, an outstanding question remains; how does chloroplastic ROS intersect with apoplastic ROS? We found that PAP cannot induce ROS accumulation in the chloroplasts of *rbohD* (Fig. 2B), indicating that apoplastic ROS is required for chloroplastic ROS accumulation, at least for stomatal closure. This is consistent with other findings, where chloroplasts are a site of ROS accumulation in guard cells in response to ozone which mimics apoplastic ROS (Joo et al., 2005).

It is unclear how PAP, ABA or apoplastic ROS leads to perturbation of the photosynthetic electron transport chain to generate ROS in chloroplasts, but one possible pathway is via Ca^2+^ signaling or homeostasis which can influence the activity of several PSI proteins and photoprotection mechanisms such as cyclic electron flow, thereby affecting photosynthetic ROS accumulation (Hochmal et al., 2015; Wang et al., 2019). We previously observed Ca^2+^ influx in guard cells treated with PAP (Zhao et al., 2019), although the mechanism(s) by which this occurs is unknown. The HPCA1 kinase has been identified as an apoplastic ROS sensor which activates plasma membrane calcium channels for influx of Ca^2+^ into guard cells for stomatal closure (Wu et al., 2020). However, HPCA1 likely does not function in isolation as it acts in concert with various signaling kinases such as CBL4, CIPK26 and OST1 during systemic cell-to-cell ROS and Ca^2+^ signaling (Fichman et al., 2022).

Additionally, it has been proposed that chloroplast and mitochondrial ROS may be interconnected through the “malate valve”, where production of excess chloroplast ROS through the photosynthetic electron transport chain is partially ameliorated via export of reducing equivalents such as malate to the mitochondria. Malate is then oxidized to NADH, with subsequent NADH oxidation leading to ROS production in mitochondria (Zhao et al., 2018, 2020). Treating guard cells with DCMU to block accumulation of chloroplast ROS resulted in no increase in H_2_DCFDA fluorescence when measured across the entire cell (Fig. 1B). By contrast, chloroplast H_2_O_2_, measured using chloroplast-targeted roGFP-Orp1, continued to increase even when mitochondrial ROS accumulation was chemically inhibited (Postiglione and Muday, 2023). These observations suggest that chloroplast ROS precedes mitochondrial ROS in stomatal closure, in agreement with the malate valve model, and could also explain why apoplastic ROS is required for both chloroplast ROS (Fig. 2) and mitochondrial ROS (Postiglione and Muday, 2023).

We observed that chloroplast ROS is in turn required for PAP-induced apoplastic ROS (Fig. S7), which raises the possibility of a ROS positive feedback loop between the organelles. One potential mechanism is linked to the possibility of chloroplast ROS export into the cytosol and nuclei directly via aquaporins (Byrt et al., 2023), or indirectly via mitochondria as described above. Regardless of the mechanism(s) by which chloroplast ROS is relayed to the cytosol, there are multiple cytosolic and/or nuclear-localized redox sensitive kinases such as OXI1 and MAP kinases (MPKs) which can activate RBOH proteins (Jalmi and Sinha, 2015).

### CPKs transcriptionally induced by PAP integrate apoplastic ROS and ion channel regulation

Interestingly, some of the PAP-induced CPKs have multiple functions in guard cells i.e., activating both RBOHD and SLAC1 (Table S1). Notably, the PAP-induced CPKs activate SLAC1 through at least two phosphorylation sites, S59 and S120 (Fig. 3C, S10) unlike the previously reported specificity of other CPKs for S59 and OST1 for S120 (Brandt et al., 2015). The multiplicity in target proteins and phosphosites of PAP-induced CPKs may translate to an additive effect for stomatal regulation, since the CPKs identified to have multiple targets (i.e., CPK13, CPK32 and CPK34) provided the greatest degree of complementation in preventing leaf water loss when expressed in the *ost1* mutant background (Fig. 3E). The upregulation of these different CPKs that are distinct to those involved in ABA signaling identifies PAP as a node that connects the chloroplast, ROS and calcium signaling with ion channel activity for stomatal regulation.

Our finding that the induction of stomatal closure by PAP occurs via RBOHD but not RBOHF, while ABA acts primarily via RBOHF (Fig. 2) (Kwak et al., 2003), provides a distinct delineation between these two drought-induced signaling molecules and their downstream pathways. Intriguingly, plants under prolonged water stress have reduced stomatal sensitivity to ABA, which is exacerbated the longer the water stress is sustained (i.e. 24 hours vs 14 days) (Peng and Weyers, 1994). During drought ABA accumulates rapidly (∼1-3 days) in *Arabidopsis* compared to the slower buildup of PAP (∼4-9 days) (Estavillo et al., 2011). This temporal separation of ABA and PAP accumulation may be relevant. It could lead to ABA activating particular CPKs and OST1 to produce ROS, Ca^2+^ and activate SLAC1 to close stomata in the earlier stages of drought. With prolonged drought stress, subsequent PAP accumulation activates the parallel CPK-RBOHD-SLAC1 pathways described herein to maintain stomatal closure. Taken together, this suggests that chloroplasts, via signals such as PAP, co-regulate the same proteins targeted by ABA to provide a time-gated reinforcement mechanism for stomatal regulation.

### Is chloroplast signaling conserved across cell types?

Our observations align with the hypothesis of cell type-specific functionalization of chloroplasts and the model of the sensory plastid, which proposes specialized environmental-sensing roles for chloroplasts that differ by cell type (Mackenzie and Mullineaux, 2022). PAP contributes to a reduction in ROS in vascular bundles and mesophyll cells, which is consistent with PAP being a chloroplast ROS-triggered signal in foliar tissue (Chan et al., 2016a) that leads to the quenching of excess ROS via up-regulation of the H_2_O_2_ scavenger *APX2* in this tissue type (Estavillo et al., 2011). In contrast, PAP induces the induction of chloroplast PSI ROS in guard cells (Fig. 1). Guard cells obtain energy and carbohydrates via different routes from mesophyll chloroplasts for stomatal opening regulation, even though chloroplasts from both cell types are photosynthetically competent (Lawson et al., 2008, 2003; Lim et al., 2022). Guard cells also have a higher proportion of chloroplasts associated with the nucleus (Fig. S5) compared to mesophyll cells, where direct H_2_O_2_ transfer from chloroplasts to the nucleus was reported (Exposito-Rodriguez et al., 2017). Given that SAL1 is expressed in multiple leaf cell types but is concentrated in the vascular bundles (Estavillo et al., 2011), it may be that PAP accumulates, and functions, differently in different cell types during stress. We summarize these putative cell-specific differences in a proposed model (Fig. 5).

**Figure 5.**
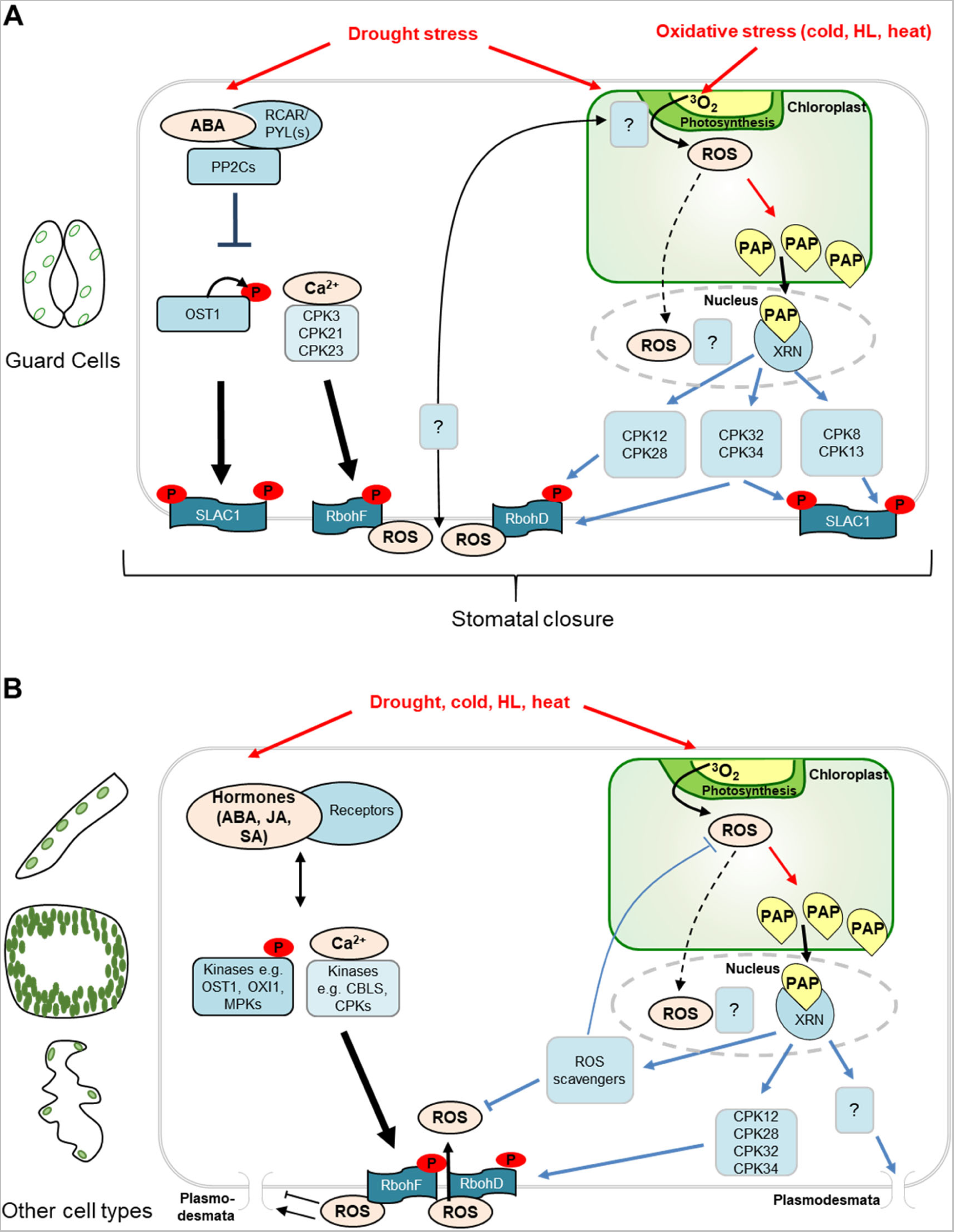
Putative model for differential interactions between PAP and ROS in specialized cell types. (A) In guard cells, PAP functions in parallel to ABA signaling to induce ROS production in chloroplasts, the nucleus, and the apoplast. Some of the CPKs transcriptionally induced by PAP can activate RBOHD for apoplastic ROS production, in contrast to ABA signaling via OST1 and other CPKs which target RBOHF. Some of the PAP-induced CPKs can activate both RBOHD and SLAC1, thus explaining the ability of PAP to function in parallel to ABA. Both chloroplast and apoplastic ROS are required for stomatal closure, and ROS production in both compartments are interdependent via unknown mechanisms. (B) In other leaf cell types such as bundle sheath, mesophyll and epidermal pavement cells, PAP has different effects on ROS accumulation and processes downstream of ROS. While PAP-induced CPKs may activate RBOHD for apoplastic ROS production and regulate plasmodesmal opening, PAP also induces expression of ROS scavengers which suppress ROS accumulation in these cell types. There is no evidence that PAP induces chloroplast or nuclear ROS in cell types other than guard cells. Other hormones and Ca^2+^ signaling can also activate RBOH-mediated apoplastic ROS production, but their intersection with PAP in these cell types remains unclear.

Our preliminary observations suggest that PAP-mediated chloroplast signaling can intersect with ROS and Ca^2+^ across different cell types and cellular processes (Fig. 4). In some cases, the mechanisms by which these pathways intersect are likely similar to that in guard cells, while in others there may be differences that in part reflect the roles of PAP in different cell types. The enhanced extracellular ROS production when challenged with flg22 in *sal1* (with constitutively accumulated PAP) (Fig. 4A) may be explained by PAP-upregulated CPKs accentuating the RBOHD-mediated ROS burst in epidermal pavement cells directly exposed to flg22 in the assay medium. In contrast, a steady state of increased PAP levels with coinciding lower vascular ROS (Estavillo et al., 2011) and no additional stress elicitation might explain the increased plasmodesmal permeability observed (Fig. 4C) as different ROS levels can modulate plasmodesmal aperture (Cui and Lee, 2016; Fichman et al., 2021).

Our results herein contribute towards resolving enigmas pertaining to cell specialization and sensory plastids. PAP induces accumulation of both chloroplastic and apoplastic ROS; with both subcellular sources of ROS being necessary for stomatal closure (Fig. 1–2). We establish the epistatic hierarchy of signaling across the cell, with PAP-induced transcriptional up-regulation of CPKs enabling activation of RBOHD for the apoplastic ROS burst, which functions in tandem with the chloroplastic ROS generation and stomatal closure (Fig. 3). The PAP-induced CPKs can target both RBOHD and the anion channel SLAC1, including a SLAC1 phosphosite previously considered to be OST1-specific, thus enabling bypass of the ABA-OST1 pathway for stomatal closure (Fig. 3). Finally, the effect of PAP extends beyond guard cells as it intersects with ROS in diverse cell types and processes (Fig. 4). In conclusion we illustrate how chloroplast retrograde signals can intersect with and influence multiple signaling pathways to regulate plant processes in a cell type-dependent manner.

## Materials and Methods

Extended Methods and Materials are available in *SI Appendix*.

### Plant material

*Arabidopsis* wild type used in this study is Columbia-0 (Col-0). Mutants and transgenic lines are in the Col-0 background except for *ost1*-2, which is in the Landsberg erecta background. Lines used in this study are *sal1*-6 (SALK_020882(Estavillo et al., 2011)); *sal1*-8 (CS66977; (Estavillo et al., 2011)); tAPX OE (Murgia et al., 2004); *ost1*-2 (*ost1*-2,(Mustilli et al., 2002)); *rbohD* (CS9555, (Torres et al., 2002)); *rbohF* (CS9557, (Torres et al., 2002)); *rbohDF* (CS9558, (Torres et al., 2002)); *slac1*-4 (SALK_137265, (Vahisalu et al., 2008)); *slac1-*S59A/S120A (SLAC1 S59A/S120A; (Brandt et al., 2015)), *ost1* (SALK_008068, (Yoshida et al., 2002)). Candidate CPKs were transformed in the Col-0 *ost1* mutant using the agrobacterium transformation floral dip method (Clough and Bent, 1998) and hygromycin screen (Harrison et al., 2006), for candidates cloned in the pMDC32 binary vector (Curtis and Grossniklaus, 2003) with the cauliflower mosaic virus 35S promoter replaced with the guard cell promoter pGC1 (Yang et al., 2008). For stomatal aperture, flg22 ROS and water-loss assays, *Arabidopsis* plants were grown on soil with 16 h light at 22 °C, except for the *rboh* mutants and the respective Col-0 control, which were grown at 12 h light. For confocal microscopy and ozone treatment assays, *Arabidopsis* plants were grown on soil in 12 h light. For bombardment assays, *Arabidopsis* plants were grown on soil with 10 h light at 22 °C.

### Microprojectile bombardment assays

Bombardment assays of cytosolic GFP endoplasmic reticulum RFP (red fluorescent protein) were performed as described in Tee et al. (2022). Bombardment sites were imaged 16 hours after bombardment using a confocal microscope (LSM Zeiss 800) with a 20× water dipping objective (W N-Achroplan 20×/0.5). GFP was excited with a 488 nm argon laser and collected at 505-530 nm, while RFP was excited with a 561 nm DPSS laser and collected at 600-640 nm.

### Flg22-induced chemiluminescence ROS assay

Chemiluminescence ROS assays were performed as described in Albert et al. (2015). See extended methods.

### Ozone treatment

Three-week-old *Arabidopsis* plants were exposed to 50 ppb ozone for six hours from 9:00 to 15:00. Plants were then transferred back into the growth chamber, with damage visually being scored and photographed 24 hours post exposure. For electrolyte leakage measurements, whole rosettes were collected after the end of the ozone exposure (both ozone treated and those in the control condition, i.e., plants left in the growth chamber at <20 ppb ozone) and placed into 20 mL of MilliQ H_2_O. Samples were measured using a Mettler conductivity meter (Model FE30) to obtain ion readings 24 hours post the initial ozone exposure. Samples were frozen completely, defrosted, and then measurements were taken again to obtain total ion content, with values plotted as a % total ion lost.

### Stomatal aperture assay

Stomatal aperture assays were performed as per Pornsiriwong et al. (2017). See extended methods.

### Microscopy for epidermal peels

Quantification of ROS in guard cells of epidermal peels was measured using 20 µM 2’,7’-dichlorodifluorescein diacetate (H_2_DCFDA; Invitrogen) according to Pornsiriwong et al. (2017), with epidermal peels treated as per Pornsiriwong et al. (2017). Fluorescence images were taken using a Leica inverted confocal microscope using a 496-nm excitation line of an argon multiline laser. H_2_DCFDA fluorescence emission was detected at 505-525 nm, while chloroplast auto-fluorescence was detected at 680-700 nm. Images were quantified using Fiji (NIH), with relative change in fluorescence of ROS determined with images before and after treatment.

Visualization of intracellular and apoplastic ROS in guard cells was performed using 20 µM H_2_DCFDA or 100 µM peroxy orange 1 for H_2_O_2_ (PO1; Merck Sigma-Aldrich), and 100 µg/ml Oxyburst Green H2HFF-BSA (Thermo Scientific), respectively. Markers for chloroplasts and nuclei were chlorophyll fluorescence and 500 µM 4′,6-Diamidino-2-phenylindole dihydrochloride, 2-(4-Amidinophenyl)-6-indolecarbamidine dihydrochloride (DAPI dihydrochloride; Merck Sigma-Aldrich), respectively. Qualitative visualization experiments were performed using a Leica DM5500 epifluorescence microscope, with appropriate filter cubes for the respective fluorescent markers.

### HEK293T RBOHD ROS assay

A prepared 96-well plate with HEK293T cells transfected with genes of interest [see extended methods for cloning specifics, transfection protocol, and protein extraction and western blot associated, adapted from Kimura et al. (2022)] was gently washed with 100 µL Hank’s Balanced Salt Solution [HBSS, with Ca^2+^ and Mg^+^; ThermoFisher Scientific], and then 100 µL of the assay buffer [29.9 mL HBSS, with Ca^2+^ and Mg^+^, 90 µL of 0.002 g horseradish peroxidase dissolved in 100 µL HBSS solution, 18.8 µL of 0.4 M luminol] was added to each well. The plate was put into a plate reader capable of polarization mode dispersion to measure luminescence at 37 °C. First, luminescence was measured for 15 minutes for cells only in the assay buffer [measurement time 1 s/well, interval 1 min/well], then 50 µL of 3 µM ionomycin solution dissolved in the assay buffer was dispensed in each well and measured for another 15 minutes.

### Electrophysiology

SLAC1-mediated anion currents in oocytes were performed as per Pornsiriwong et al. (2017). See extended methods.

### Statistical analyses

Data presented as boxplots have the middle line marking the median, the box indicating the upper and lower quartiles, and the whiskers showing the minimum and maximum values within 1.5× interquartile range. Data were analyzed using RStudio 2021.09.01 Build 351/R version 4.0.3. See extended methods for specifics.

## Acknowledgments

We thank Professor Julian Schroeder for kindly providing the seeds of *slac1-S59A/S120A*. We thank the Western Sydney University Confocal Bio-Imaging Facility for usage of the confocal microscope. We thank Professor Wah Soon Chow (ANU) for guidance on usage of the DCMU inhibitor, and Laura Bailey for assistance with the DAPI and Oxyburst Green H2FF-BSA experiments. We received financial support from the ARC Laureate Fellowship (FL190100056 to BJP); ANU scholarships (EET); ARC Discovery Early Career Researcher Award (DE210100466 to KXC); ARC Future Fellowship (FT210100366 to ZHC); European Research Council (grant agreement 725459, “INTERCELLAR”) and the Biotechnology and Biological Research Council (Institute Strategic Programme ‘Plant Health’ BBS/E/J/000PR9796) (AB and CF), Australian Endeavour Award (6975_2018) and CSIRO Synthetic Biology Future Science Platform (MC), and the Academy of Finland (decisions #275632, #28313, #312498 and #323917 for MW).

