## Supplementary Methods and Data for "Cell-specialized chloroplast signaling orchestrates photosynthetic and extracellular reactive oxygen species for stress responses"

**This PDF file includes:**

Extended Material and Methods

Figures S1 to S14

Tables S1 and S3

SI References

### Extended Material and Methods

*Stomatal aperture assay and measurements*

For any epidermal peel assay, samples were harvested from three to five-week-old plants, with a minimum of three biological replicates used per treatment and genotype. The epidermal layer was gently adhered adaxial side down to the bottom of the experimental chamber (i.e., small petri dish) coated with optically clear and pressure-sensitive silicon adhesive (Factor II Inc.). Samples were incubated in Opening Buffer [50 mM KCl, 5 mM MES titrated to pH 6.1 with NaOH] for two hours, and then transferred to an inverted bright-field microscope capable of 400× magnification. An initial image was taken and then another after 10 minutes to ensure the stomata were open before subsequent assays. Samples were then washed gently three times with Measuring Buffer [10 mM KCl, 5 mM MES titrated to pH 6.1 with $Ca(OH)_2$], and then treated with the appropriate chemical dissolved in Measuring Buffer with images taken every 10 minutes. Stomatal aperture data are expressed as relative change over the control period (i.e., change from the initial image taken before washes and treatment) after 20 minutes. The stomatal pore area was determined using Fiji (NIH) where pore areas were converted to pixels, and relative change in pixels per pore area between images were compared.

*flg22-induced chemiluminescence ROS assay*

The following protocol was adapted from Albert et al. (2015). A 96-well flat bottom white plate was prepared with 120 µL MilliQ $H_2O$ per well per sample. *Arabidopsis* leaf discs were collected and placed with the abaxial side down and left to recover overnight. Leaves were placed in a spatially optimized design and treatment added in a random order to reduce position effects, factored into the statistical analyses. Water was removed the next day and a mix of 90 µL MilliQ $H_2O$ + 10 µL luminol master mix [200 µM luminol L-012 (Wako Chemicals, United States) and 10 µg/mL horseradish peroxidase] was added to each well. An infinite M1000Pro TECAN plate reader (Tecan Group Ltd, Switzerland) was used to measure luminescence with the following settings: Shaking (Orbital) Duration: 3 s; Shaking (Orbital) Amplitude: 1 mm; Shaking (Orbital) Frequency: 582 rpm;

Interval Time: Minimal; Mode: Luminescence; Attenuation: NONE; Integration Time: 100ms; Settle Time: 1000ms. Measurements were taken for approximately 10 minutes to ensure equilibration was achieved, before treatments (mock or flg22 added for a total concentration of 500 nM per well) were added.

*HEK293T cloning vectors and transfection*

CPK genes were cloned into pEF1-MCS-3Myc, an altered pEF1/*myc*-His vector (Invitrogen)(Kimura et al., 2020). The vector with RBOHD, pcDNA3.1-3FLAG-RBOHD, was previously described (Kaya et al., 2019). A midiprep for DNA used for transfecting HEK293T plasmids was prepared using the NucleoBond® Xtra Midi Plus Kit (MACHEREY-NAGAL GmbH & Co). HEK293T cells were sub-cultured for four weeks, with transfection and ROS assays performed on cells two to four weeks of age. A cell suspension was adjusted to a cell density of 1.0-2.0 × $10^5$ cells/mL, then 130 µL was aliquoted into each well of a white Corning® BioCoat® Poly-D-lysine coated 96 well plate, then incubated at 37 °C 5% $CO_2$ for 12-24 hours. A plasmid ratio of 5:1 co-transfection ratio of pcDNA3.1 vector:pEF1/*myc*-His vector was used (100 ng to 20 ng). For each combination tested, 12 wells were transfected with the exception of the non-transfection control. Plasmid DNA was diluted to be 100 ng/µL so total DNA content added was 1.2 µL per well. 6.3 µL Opti-MEM (ThermoFisher Scientific) was transferred into sterile microcentrifuge tubes, and 0.36 µL per well to be transfected of GeneJuice® Transfection Reagent (Novagen®, Millipore) was added directly to the Opti-MEM, mixed thoroughly by vortexing. The mixture was incubated at room temperature for five minutes, then plasmid DNA was added, mixed by gently pipetting and incubated further for another five minutes. The mixture was added dropwise to prepared cells, and the whole plate was gently rocked to ensure even distribution. Cells were then incubated for another 48 hours at 37 °C 5% $CO_2$. CPK candidates were screened either in combination with the pcDNA3.1 empty vector of RBOHD for a minimum of three separate plates. Data per plate was from three specific wells, with each plate designed so there were three replicates per transfection combination, and an average measured between the three.

*Protein extraction and western blot for HEK293T cells*

The medium was removed from three selected wells from each transfection combination. 50 µL of protein sampling buffer [50 mM Tris-HCl, 2% SDS, 10% glycerol, 10% 2-mercaptoethanol, 300 mM DTT] was added and left to incubate for five minutes. The lysates were collected into microcentrifuge tubes, and before protein loading, samples were pipette syringed to fragment genomic DNA. For HEK293T cells transfected with plasmids containing the FLAG tag, protein amount loaded was 10 µL, while plasmids containing the *myc* tag, protein amount loaded was 50 µL with the exception of GFP, in which only 5 µL was loaded. A wetblot transfer onto Immobilon® Transfer Membranes (Merck Millipore) was performed, and membranes were probed with anti-FLAG (1:3000 dilution) and anti-β ACTIN (1:5000 dilution), or anti-c-Myc (1:3000 dilution) followed by IRDye 800 anti-mouse-IgG (IRDye ® 800CW Goat anti-Mouse IgG (H + L), 0.5 mg; LI-COR Biosciences).

Xenopus Laevis *surgery and oocyte preparation*

Female *Xenopus laevis* frogs were anaesthetized by submersion for 30 min in 3-Aminobenzoic acid ethyl ester (1.5 g/L) (Sigma-Aldrich), then placed on ice to slow blood flow; surgery was conducted as previously described (Fairweather et al., 2021, 2015). Ovary sac sections were maintained in $OR^{2-}$ buffer (82.5 mM NaCl, 2.5 mM KCl, 1 mM $MgCl_2$, 1 mM $Na_2HPO_4$, 5 mM HEPES-NaOH, pH 7.8) in sterile Petri dishes and cut into small clumps of ~30-50 oocytes. Digestion of folliculated oocytes into individually accessible de-folliculated oocytes was conducted by incubating cut ovary sections in 1.5 mg/mL collagenase D (Sigma-Aldrich) dissolved in $OR^{2-}$ (pH 7.8) buffer for 2 hours at 28 °C followed by 4 hours at 18 °C. The success of digestion and de-folliculation was assessed following the washing and examination of oocytes using 2 L of $OR^{2-}$ buffer followed by 2 L $OR^{2+}$ buffer ($OR^{2-}$ supplemented with 1.5 mM $CaCl_2$ and 50 µg/mL gentamycin, pH 7.8). Selected healthy oocytes were maintained at 16−18 °C in $OR^{2+}$ until required for electrophysiology experiments. Maintenance of animals and preparation of oocytes was approved by the Australian National University animal ethics review board (ANU Protocol A2017/36).

*Electrophysiology recordings*

New CPKs of interest were first cloned in pCR™8/GW/TOPO vector and subcloned into a destination pGEM-HE oocyte vector used in Pornsiriwong et al. (2017). For SLAC1 phosphosite mutations, site-directed mutagenesis was performed using the QuikChange XL Site-Directed Mutagenesis Kit (Agilent Technologies) [for SLAC1-S120A: GCAAACAAAAG**G**CTTTATTGCCTTCTAT and ATAGAAGGCAATAAAG**C**CTTTTGTTTGC; for SLAC1-S59A: GCAGACAGGTTGCGCTAGAGACAGG and CCTGTCTCTAGCG**C**AACCTGTCTGC]. Plasmids were first linearization with either restriction enzymes NheI or SbfI, followed by cRNA synthesis by mMESSAGE mMACHINE® T7 kit (Thermo Fisher Scientific). Prepared *Xenopus laevis* oocytes were injected with RNase-free $H_2O$, or cRNA. For kinase or SLAC1/SLAC1 mutant controls, 25 ng was injected. SLAC1/SLAC1 mutants were mixed with individual CPKs for a ratio of 2:1. Whole-cell currents were recorded from oocytes 3 days post injection using an Axoclamp 2B two-electrode clamp circuit. Measurements were recorded in an $I_{anion}$ measuring solution [48 mM NaCl, 48 mM CsCl, 1 mM $MgCl_2$, 1 $CaCl_2$, 10 mM MES, titrated to pH 5.6 with 5 M HCl], with the following protocol: holding at 0 mV for 1 s, testing from +50 to -130 mV for 10 × 10 s cycles (holding voltage taken every 20 mV intervals), and holding at 0 mV for 1 s. Two-electrode voltage-clamp recordings were performed with 1 × LU and 10 × MGU head stages connected to a Geneclamp 500B electronic amplifier (Axon Instruments, Union City, U.S.A). The output signal was amplified 10 times and filtered at 1 kHz, with the analogue signal converted into digital by a Digidata 1322A (Axon Instruments). Data were sampled at 10 Hz using pCLAMP software (Axon Instruments), and steady state readings were taken as an average at the end of the 10 second holding voltage step using the program Clampfit 10.7.0.3 (Molecular Devices).

*Extended statistical analyses*

For microprojectile bombardment assays, data were analyzed by bootstrap method (*medianBootstrap,* (Johnston and Faulkner, 2021)). For flg22-induced chemiluminescence ROS assays; ion leakage assay after ozone treatment; stomatal aperture assays; ROS changes in stomatal components or whole stomata; HEK293T ROS assay; and electrophysiology measurements showing SLAC1 anion currents at -130 mV; a linear mixed-effects model (with independent factors specified in figure legends) was applied using the R package, *lmerTest*. For experiments where multiple replicates were taken from a given biological replicate (*i.e.,* different stomata from a biological replicate), the random effect was 'biological replicate'. ANOVAs specified are ANOVA Satterthwaite's Method, with significant differences between factors (denoted in figure legends) determined by post hoc Tukey HSD using the R package, *emmeans.*

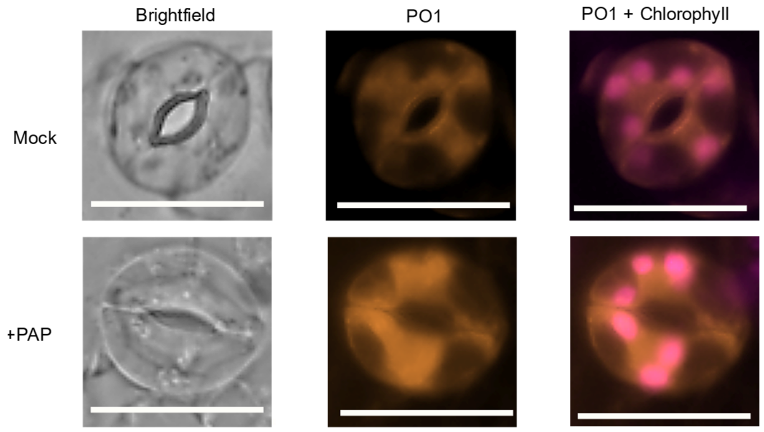


**Figure S1. PAP treatment increases $H_2O_2$-responsive peroxy orange 1 (PO1) fluorescence in guard cells**

Representative images for guard cells cotreated with 100 µM PO1 and either mock or 100 µM PAP for 30 min in the light. Each condition contains a minimum of three wild type plants with a minimum of n ≥ 10 stomata per treatment combination. Scale bar = 12 µm.

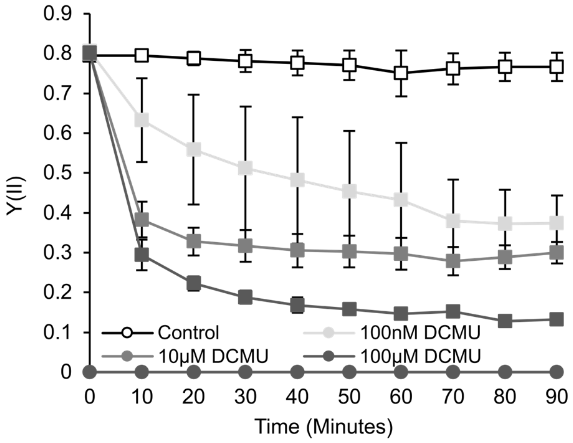


**Figure S2. Reduction of effective PSII quantum yield (Y(II)) in mesophyll and epidermal tissue by DCMU**

Reduction of Y(II) was monitored over time in response to three concentrations (100 nM, 10 μM and 100 μM) of DCMU. Mesophyll tissue is represented in squares while epidermal tissue is represented in circles. Values are averages of three replicates per treatment with error bars representing SD.

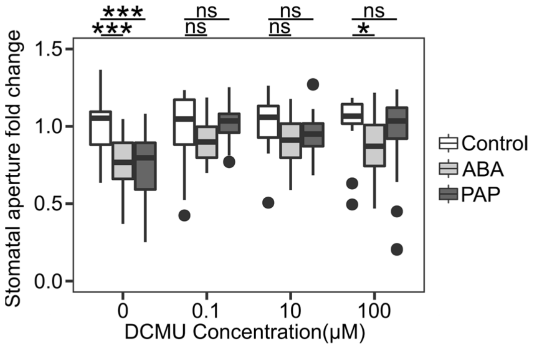


**Figure S3. PAP and ABA-mediated stomatal closure is differentially inhibited by various concentrations of DCMU**

Stomatal closure in response to 100 μM ABA or 100 μM PAP with 0, 0.1, 10 or 100 μM DCMU. Each condition contains a minimum of three wild type plants with a minimum of n ≥ 26 stomata per treatment combination. 'DCMU' and 'Treatment' significantly impacted closure with a significant interaction between the two (ANOVA; F=13.3, df=3, p<0.001; F=22.4, df=2, p<0.001; F=4.1, df=6, p<0.001). Significant differences from respective DCMU Control denoted by *, p<0.05 or ***, p<0.001.

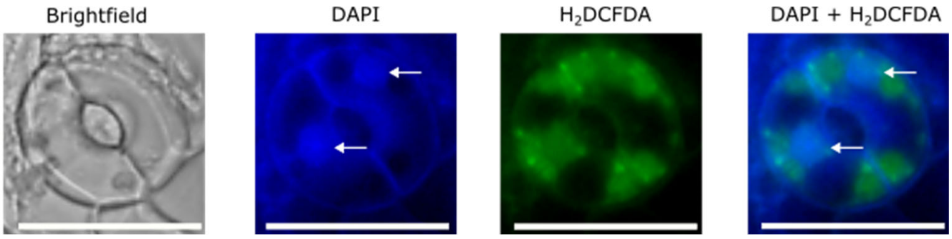


**Figure S4. PAP treatment induces a ROS burst in nuclei of guard cells**

Co-localisation of PAP-induced ROS was visualised in guard cells co-treated with 100 µM PAP, 500 µM DAPI and 20 µM $H_2$DCFDA. Nuclei are indicated by white arrows. Scale bar = 12 µm.

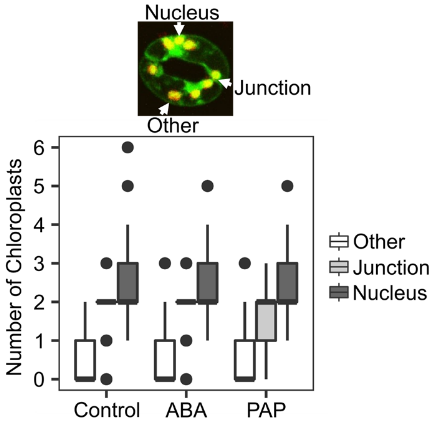


**Figure S5. Chloroplast localizations in guard cells**

Number of chloroplasts in specified localizations (indicated by white arrows) for a given guard cell. Four biological replicates with number of stomata measured ≥59 per treatment. Treatment did not have a significant effect on the chloroplast number (ANOVA; $F=0.6$, $df=2$, $p=0.6$).

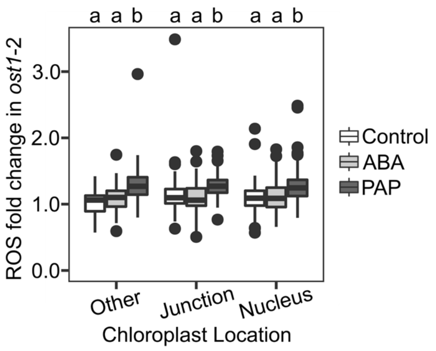


**Figure S6. Different chloroplast location types accumulate ROS similarly**

ROS fold change measurements in different chloroplast locations for a given guard cell. Three biological replicates with number of guard cells measured ≥64 per treatment. Treatment had a significant effect on ROS fold change (ANOVA; $F=76.6$, $df=2$, $p<0.001$), while chloroplast location nor the interaction between treatment and chloroplast location did not (ANOVA; $F=0.51$, $p=0.6$; $F=1.8$, $df=4$, $p=0.1$). Significant differences denoted by a and b, $p<0.01$.

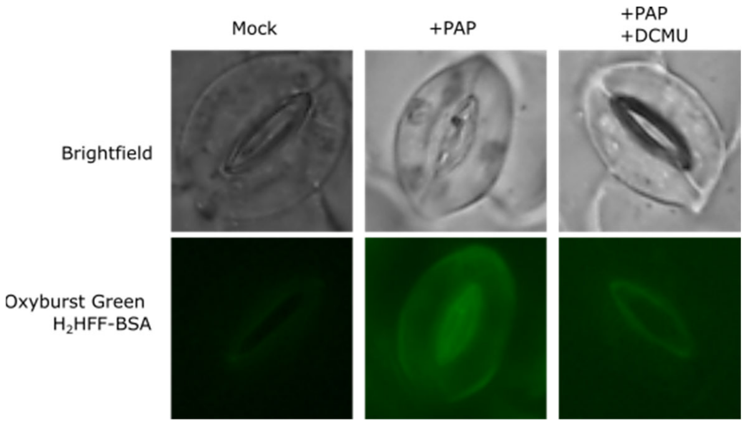


**Figure S7. Inhibition of chloroplast ROS production with DCMU diminishes PAP-induced apoplastic ROS production**

Representative images for guard cells treated with either mock, 100 µM PAP, or 100 µM PAP and 10 µM DCMU. All treatments were co-incubated in the light for 20 min with 100 µg/ml Oxyburst Green H2HFF-BSA. Each condition contains a minimum of three wild type plants with a minimum of n ≥ 20 stomata per treatment combination.

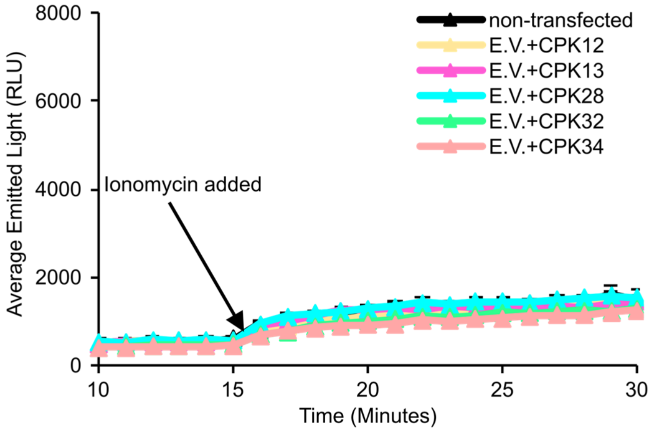


**Figure S8. Expression of CPKs alone in HEK293T cells does not result in ROS production**

ROS production of HEK293T cells transiently expressing CPKs with empty vector (E.V.). After 15 min 1 μM ionomycin was added to the medium. Values represent mean ± SEM of n = 3. The experiment was repeated in three independent experiments with similar results.

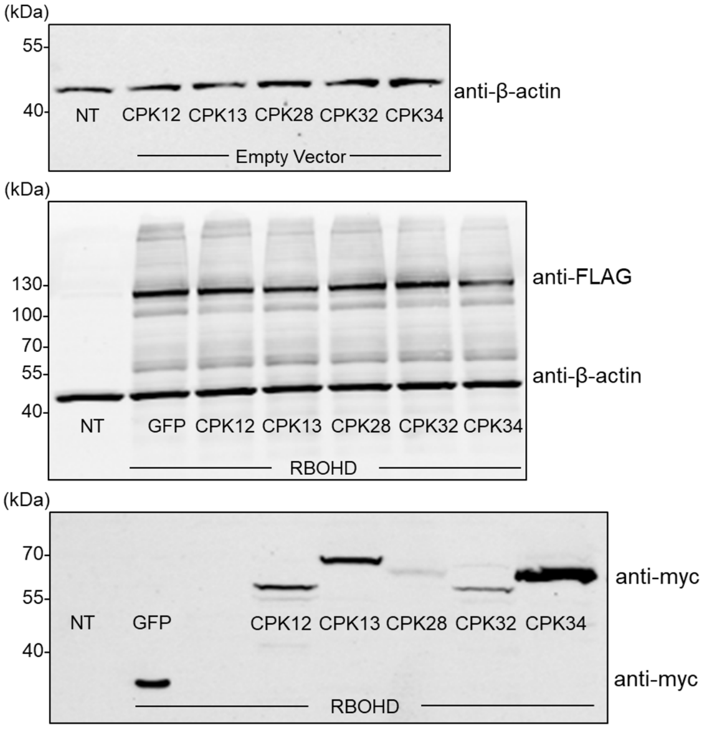


**Figure S9. Expression of CPKs and RBOHD in HEK293T cells**

Example western blots showing equal protein loading via anti-β-actin. NT stands for non-transfected control. Top figure shows equal loading associated with Fig. S8, with equal loading of total protein. Middle figure indicates equal expression of RBOHD (anti-FLAG) and equal loading of total protein (anti-β-actin), associated with Fig. 3A. Bottom figure shows proteins (either GFP as the negative control, or the CPKs) tagged with anti-myc, associated with Fig. 3A. There was unequal protein expression of the CPKs as indicated by anti-myc despite equal total protein loading.

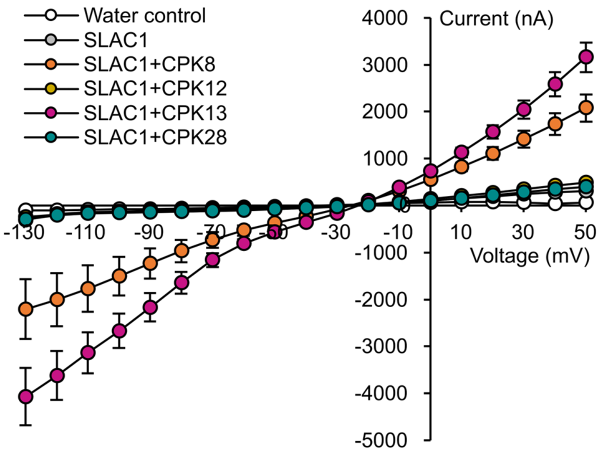


**Figure S10. Current traces of CPK candidates**

Current traces of CPK8, CPK12, CPK13 and CPK28 activations of SLAC1 anion channel in oocytes. Oocytes were co-injected with the kinase and the channel SLAC1. Values are means of four to eight oocytes ± SEM. Anion channel activity shown as trace data at voltage pulses ranging from +50 to -130 mV in 10mV decrements.

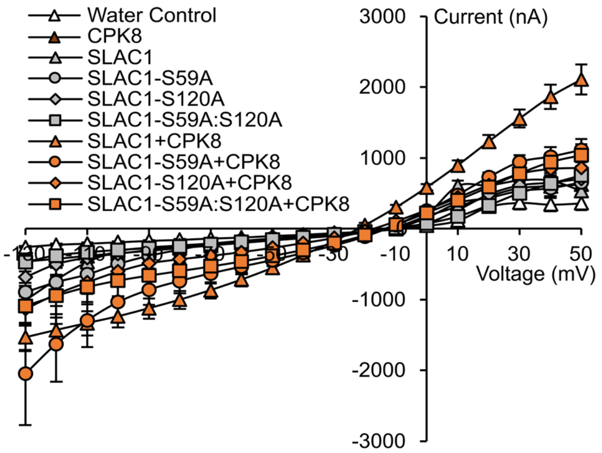


**Figure S11. Representative current traces of CPK8 with different mutated phosphosite SLAC1**

Current trace of CPK8 with mutated SLAC1 anion channel in oocytes. Oocytes were injected with CPK8 alone, the channel SLAC1 (wild type or mutated), $H_2O$ (Water control), or in combination. Values are means of a minimum of seven oocytes per combination ± SEM from a single surgery and is representative of two independent repeats. Anion channel activity shown as trace data at voltage pulses ranging from +50 to -130 mV in 10 mV decrements.

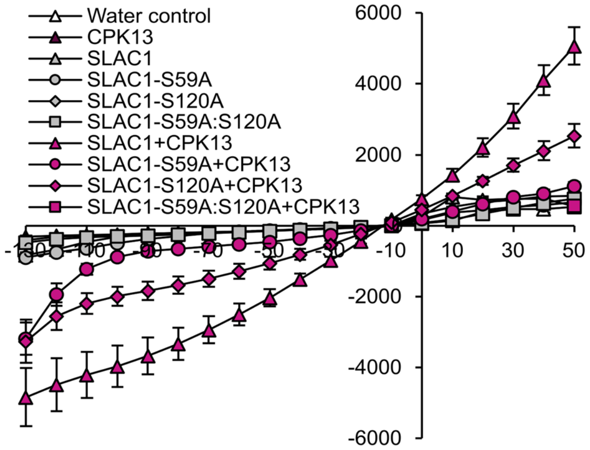


**Figure S12. Representative current traces of CPK13 with different mutated phosphosite SLAC1**

Current trace of CPK13 with mutated SLAC1 anion channel in oocytes. Oocytes were injected with CPK13 alone, the channel SLAC1 (wild type or mutated), $H_2O$ (Water control), or in combination. Values are means of a minimum of six oocytes per combination ± SEM. Anion channel activity shown as trace data at voltage pulses ranging from +50 to -130 mV in 10 mV decrements.

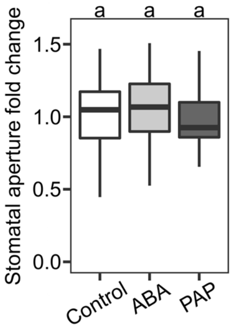


**Fig. S13 *slac1*-4 mutants are unresponsive to ABA and PAP**

Stomatal closure of *slac1*-4 mutants in response to 100 μM ABA or 100 μM PAP. Each treatment contains five biological replicates with a minimum of n > 36 stomata per treatment. 'Treatment' did not impact closure (ANOVA; F=1.3, df=2, p=0.3).

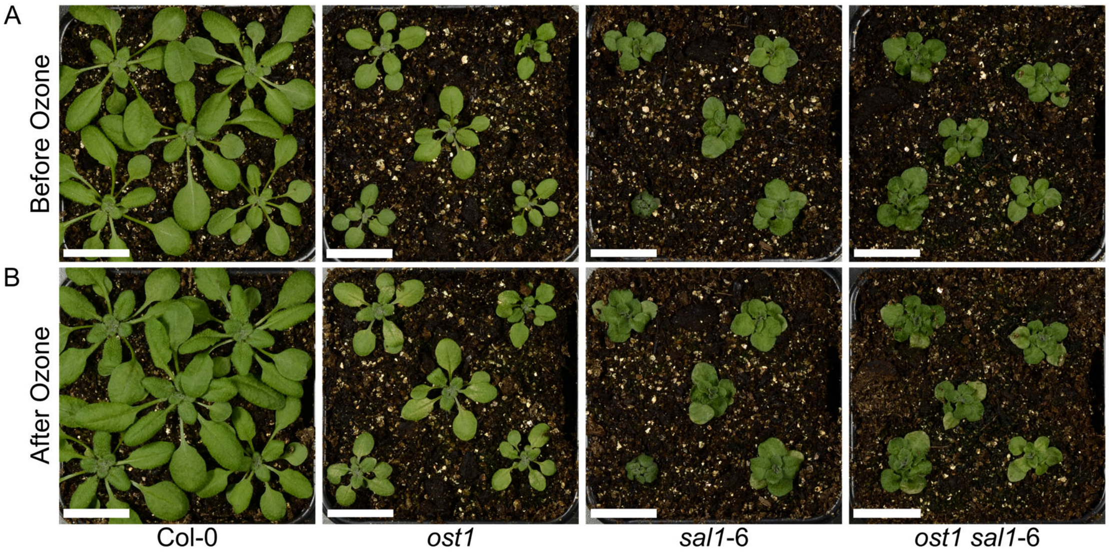


**Figure S14. Ozone treatment on *sal1*-6 mutants**

(A) Before and (B) after pictures of 3-week-old plants, taken 24 hours after the end of a six-hour exposure to 350 ppb ozone (i.e., 30 hours post initial exposure). Scale bar = 2 cm.

**Table S1. Summary of PAP-transcriptionally induced candidate function in RBOHD and SLAC1 activity, and water-loss assay in *ost1* background**

| Gene | RBOHD | SLAC1 | Complementation of *ost1* |
|---|---|---|---|
| CPK8 | n/a | Yes | Minor |
| CPK12 | Yes | No | Minor |
| CPK13 | No | Yes | Partial |
| CPK28 | Yes | No | n/a |
| CPK32 | Yes | Yes | Partial |
| CPK34 | Yes | Yes | Partial |

**Table S3. Microarray analysis of altered gene expression in *sal1*-8 relative to Col-0 as per Estavillo et al. (2011).** Changes were considered significant at an FDR correction level of PPDE(>P) >0.95 and a fold change greater than 1.5 fold.

| Locus Identifier | Annotation | Fold Change | Bayes.p | Reference |
|---|---|---|---|---|
| AT1G05570 | ATGSL06/CalS1 | -1.66 | 0.03 | (Cui and Lee, 2016) |
| AT1G22610 | MCTP6 | 1.77 | 7.26E-05 | (Brault et al., 2019) |
| AT2G17120 | LYM2 | -1.78 | 0.01 | (Faulkner et al., 2013) |
| AT2G23770 | LYK4 | -1.97 | 0.01 | (Cheval et al., 2020) |
| AT3G45600 | TET3 | -1.50 | 0.01 | (Fernandez-Calvino et al., 2011) |
| AT3G50770 | CML41 | 2.91 | 0.02 | (Xu et al., 2017) |
| AT3G57880 | MCTP3 | 4.12 | 0.001 | (Brault et al., 2019) |
| AT3G60720 | PDLP8 | -1.66 | 0.02 | (Ye et al., 2017) |
| AT4G03550 | ATGSL05 / CalS12 | -1.53 | 0.01 | (Nishimura et al., 2003) |
| AT5G46330 | FLS2 | 2.55 | 1.38E-05 | (Faulkner et al., 2013) |
